# Postnatal developmental trajectory of sex-biased gene expression in the mouse pituitary gland

**DOI:** 10.1101/2022.01.05.475069

**Authors:** Huayun Hou, Cadia Chan, Kyoko E. Yuki, Dustin Sokolowski, Anna Roy, Rihao Qu, Liis Uusküla-Reimand, Mariela Faykoo-Martinez, Matt Hudson, Christina Corre, Anna Goldenberg, Zhaolei Zhang, Mark R. Palmert, Michael D. Wilson

**Affiliations:** Genetics and Genome Biology, SickKids Research Institute, Toronto, ON, Canada; Department of Molecular Genetics, University of Toronto, Toronto, ON, Canada; Donnelly Centre for Cellular & Biomolecular Research, Toronto, ON, Canada; Interdepartmental Program of Computational Biology and Bioinformatics, Yale University, New Haven, CT, USA; Department of Pathology, Yale School of Medicine, New Haven, CT, USA; Department of Cell and Systems Biology, University of Toronto, Toronto, ON, Canada; Department of Computer Science, University of Toronto, Toronto, ON, Canada; Division of Endocrinology, The Hospital for Sick Children, Toronto, ON, Canada; Institute of Medical Science, University of Toronto, Toronto, ON, Canada; Departments of Pediatrics and Physiology, University of Toronto, Toronto, ON, Canada

## Abstract

The pituitary gland regulates essential physiological processes such as growth, pubertal onset, stress response, metabolism, reproduction, and lactation. While sex biases in these functions and hormone production have been described, the underlying identity, temporal deployment, and cell-type specificity of sex-biased pituitary gene regulatory networks are not fully understood. To capture sex differences in pituitary gene regulation dynamics during postnatal development, we performed 3’ untranslated region sequencing and small RNA sequencing to ascertain gene and microRNA expression respectively across five postnatal ages (postnatal days 12, 22, 27, 32, 37) that span the pubertal transition in female and male C57BL/6J mouse pituitaries (n=5-6 biological replicates for each sex at each age). We observed over 900 instances of sex-biased gene expression and 17 sex-biased microRNAs, with the majority of sex differences occurring with puberty. Using miRNA-gene target interaction databases, we identified 18 sex-biased genes that were putative targets of 5 sex-biased microRNAs. In addition, by combining our bulk RNA-seq with publicly available male and female mouse pituitary single-nuclei RNA-seq data, we obtained evidence that cell-type proportion sex differences exist prior to puberty and persist post-puberty for three major hormone-producing cell types: somatotropes, lactotropes, and gonadotropes. Finally, we predicted sex-biased genes in these three pituitary cell types after accounting for cell-type proportion differences between sexes. Our study reveals the identity and postnatal developmental trajectory of sex-biased gene expression in the mouse pituitary. This work also highlights the importance of considering sex biases in cell-type composition when understanding sex differences in the processes regulated by the pituitary gland.

**Highlights:**

- Male and female mouse pituitary gland gene and miRNA expression was profiled across five postnatal ages spanning pubertal development
- Abundant sex differences in pituitary gene expression exist prior to puberty and become more prominent upon puberty
- Combining expression data from genes and miRNAs revealed 18 putative sex-biased gene targets of 5 sex-biased miRNAs
- Sex differences in the proportions of somatotropes, lactotropes, and gonadotropes are predicted to occur prior to puberty

## Background

The pituitary gland plays a central role in regulating growth, lactation, reproduction, metabolism, stress responses, and puberty. These physiological processes are mediated by hormones released from five main anterior pituitary cell-types: growth hormone (GH) from somatotropes, prolactin (PRL) from lactotropes, follicle-stimulating hormone (FSH) and luteinizing hormone (LH) from gonadotropes, thyroid-stimulating hormone (TSH) from thyrotropes, and adrenocorticotrophic hormone (ACTH) from corticotropes (Ooi et al., 2004). Unlike the more glandular anterior pituitary, the more neural posterior pituitary consists primarily of astroglial-like pituicytes as well as axonal projections from the hypothalamus which store and release oxytocin and vasopressin into the systemic circulation (Baylis and Ball, 2000; Bucy, 1930). Non-hormone producing pituitary cells, including stem cells, and folliculostellate cells, are also present in the pituitary gland and support hormone-producing cells by functioning as progenitor cells and facilitating intercellular signaling within the pituitary (Fauquier et al., 2002; Yoshida et al., 2011).

Both pituitary hormone production and many physiological processes regulated by the pituitary gland are sex-biased. For example, GH is secreted in a sexually different pattern in rodents - more pulsatile in males compared to females, and regulates sex-biased gene expression in liver (Waxman and O’Connor, 2006). Moreover, the clinical presentation and prevalence of pituitary-related disorders can also differ between sexes. For example, the prevalence of prolactinoma is significantly higher in women, but men are more likely to present with macroadenomas (Agustsson et al., 2015; Mindermann and Wilson, 1994). While the sex differences in pituitary function and disease are well known, the gene regulatory networks underlying these differences remain elusive.

Several studies in rodents and humans have clearly highlighted sex differences in pituitary gland gene regulation. These studies include: targeted qPCR profiling of genes encoding for the main pituitary hormones in rat anterior pituitaries (Bjelobaba et al., 2015); serial analysis of gene expression (SAGE) in whole adult mouse pituitaries (Nishida et al., 2005a); and RNA-sequencing of adult human pituitaries as part of the Genotype-Tissue Expression (GTEx) project (Gershoni and Pietrokovski, 2017; Lopes-Ramos et al., 2020; Oliva et al., 2020). Most recently single-cell RNA-seq (scRNA-seq) has been performed in male and female adult mouse and rat pituitary glands revealing genes with sex-biased expression within specific cell-types (Fletcher et al., 2019; Ho et al., 2020; Ruf-Zamojski et al., 2021). While most gene expression studies have focussed on the adult pituitary, sex differences in pre-pubertal gene expression have been revealed using qPCR in mouse and rat pituitary glands (Bjelobaba et al., 2015; Hou et al., 2017) and by RNA-seq in juvenile mouse gonadotropes (Qiao et al., 2016). While these studies suggest that some sex differences in pituitary gene regulation are established prior to puberty we still lack a comprehensive view of pituitary gene regulation during postnatal development.

Another essential aspect of gene regulation is post-transcriptional regulation by microRNAs (miRNAs). While miRNA expression has been explored in pituitary glands of several mammalian species, these studies were not focused on postnatal development in males and females (Bak et al., 2008; Bottoni et al., 2007; Ye et al., 2018, 2015; Yuan et al., 2015; Zhang et al., 2018). While no evidence implicating miRNAs in sex-biased gene regulation in the pituitary has been reported, there are clear examples reported in the neonatal hypothalamus and pubertal liver (Hao and Waxman, 2018; Morgan and Bale, 2017).

The objective of this study was to characterize male and female pituitary gene expression during postnatal development and investigate the role of miRNAs in regulating pituitary sex differences. To achieve this, we profiled gene (3’ untranslated region sequencing, 3’UTR-seq) and miRNA (small RNA sequencing, sRNA-seq) expression in male and female mice at multiple postnatal days spanning pubertal transition to identify genes and miRNAs exhibiting known or novel sex differences. The resulting temporal gene and miRNA expression data from this study can be queried and visualized at https://wilsonlab-sickkids-uoft.shinyapps.io/pituitary_gene_mirna_shiny/. Lastly, by combining this data with published single-cell RNA-seq (scRNA-seq) datasets, we provide evidence that sex differences in cell-type proportions emerge prior to the onset of puberty and likely contribute to sex-biases in bulk gene expression.

## Methods

### Animal and tissue collection

All studies and procedures were approved by the Toronto Centre for Phenogenomics (TCP) Animal Care Committee (AUP 09-08-0097) (see **Ethics approval** for details). Conditions in which C57BL/6J mice were maintained, sacrificed, and dissected are described previously in (Hou et al., 2017).

Physical markers of puberty, vaginal opening (VO) and preputial separation (PS), were assessed everyday after weaning to determine the pubertal stage of our female and male mice respectively (Danilovich et al., 1999; Korenbrot et al., 1977; Sánchez-Garrido et al., 2013). For assessing VO, the mouse was held by her base and a sterile pipette tip was used to brush the fur and assess the opening of the vagina. For assessing PS, the mouse was held on the hopper by his base and the degree of separation was assessed by gently pushing the prepuce using a sterile pipette tip (Korenbrot et al., 1977). Upon dissection, the pituitary gland was directly moved to RNAlater (containing 10% w/v sodium citrate tribasic dihydrate and 60% w/v ammonium sulphate) following dissection and stored at −20°C until RNA extraction.

### RNA extraction

Prior to RNA extraction, pituitary tissue samples were placed into bead mill tubes containing six 1.4-mm ceramic beads (MoBio Laboratories) and homogenized for 30 s at 6.5 m/s at 4 °C using an Omni Bead Ruptor 24 bead mill. RNA was extracted with the NucleoSpin® miRNA kit (Macherey Nagel) in combination with TRIzol lysis (Invitrogen) following the manufacturers’ protocols, allowing for collection of small RNA (< 200 nt) and large RNA (> 200 nt) simultaneously into separate tubes from total RNA. RNA quantity was determined using Nanodrop and the quality was assessed by an Agilent 2100 Bioanalyzer.

### Library preparation and sequencing

To construct RNA-seq libraries, we established an automated 3’UTR-seq (QuantSeq 3’mRNA-seq; Lexogen GmbH, Vienna) using the Agilent NGS Workstation (Agilent Technologies, Santa Clara) at The Centre for Applied Genomics (TCAG) (Toronto, Canada) as per the manufacturer’s protocol (Yuki et al., 2018). Briefly, 250 ng of RNA, from the large RNA fraction, was used to generate cDNA. cDNA was amplified with 17 PCR cycles as determined by qPCR analysis using the PCR Add-on kit (Lexogen). ERCC RNA spike-in Mix 1 was added following the manufacturer’s instructions. The resulting libraries were quantified with Qubit DNA HS (Thermo Fisher, Waltham) and fragment sizes analyzed on the Agilent Bioanalyzer using the High Sensitivity DNA assay prior to sequencing. Sequencing was performed at TCAG on the HiSeq 2500 v4 flow cell (Illumina, San Diego) with SR50 bp with cycles extended to 68bp. Small RNA-seq libraries were generated from the small RNA fraction by TCAG using NEBNext Small RNA Library Kit (New England BioLabs) according to the manufacturer’s protocol in two batches: 20 ng of RNA was used for batch 1 (replicates 1-3) and 10 ng of RNA was used for batch 2 (replicates 4-6). Sequencing was performed at TCAG on a HiSeq 2500 v4 flow cell with SR50 bp.

### mRNA sequencing reads processing

FastQC (http://www.bioinformatics.babraham.ac.uk/projects/fastqc/) was used to examine the quality of sequenced reads. Next, a customized script was used to trim both the polyAs and adaptors sequences at the end of the reads. A subset of sequencing reads obtained from 3’UTR-seq show a mixture of polyAs and sequencing adapters towards the end of the reads, which are not effectively trimmed by available read-trimming tools. We developed a trimming strategy which can identify and trim off polyA sequences embedded in the adapter sequences. Only reads longer than 36 bp after trimming were used. In addition, the first 12 nucleotides were trimmed based on the manufacturer’s recommendations. After trimming, FastQC was performed again to examine read quality. At this step, we found that ribosomal reads were overrepresented in the pituitary samples through priming by oligo(dT) binding to A-rich regions in the ribosomal RNA loci. In addition, the sequence of a brain-enriched small RNA, BC1, is also represented. Thus, reads that map to these overrepresented transcripts were removed. Trimmed and filtered reads were aligned to the genome using a splice-aware aligner, STAR (version 2.5.1b) (Dobin et al., 2013), with default settings except “-- outFilterMismatchNoverLmax 0.05” for QuantSeq. Quality control of mapped RNA-seq reads was performed using Qualimap (version 2.2.1) (Okonechnikov et al., 2016).

### PolyA site identification and gene annotation modification

GENCODE version M21 was the primary annotation used. To achieve a more comprehensive annotation of the 3’UTRs, we also incorporated the 3’UTRs annotated in RefSeq, which is obtained from the UCSC database (mm10). In addition, we also identified potential polyA (pA) sites from the data. To do this, only the 3’ most nucleotide of each read is used to build a signal track for each sample. R package “derfinder” (Collado-Torres et al., 2017) was used to identify expressed regions (ER) from these signal tracks. Specifically, an average read pile-up cutoff of 1 RPM (reads per million mapped reads) was used. ERs are annotated to gene annotations (3’UTR, 5’ UTR, exons, introns, and intergenic regions) based on GENCODE version M21, allowing for overlapping categories. ERs mapped to introns and intergenic regions are further analyzed to identify novel polyA sites. To filter for potential internal polyA priming events, sequence composition around ERs is examined. ERs with a) matches 18-mer polyAs (with up to 6 mismatches) within 150 bp downstream from the ends; b) matches 7-mer polyAs (with up to 1 mismatches) within 20 bp downstream from the ends; or c) more than 50% of the As within 20 bp downstream from the ends, are removed. In addition, ERs that overlap more than 20 bp with an annotated repeat region are also excluded. Filtered ERs are first mapped to RefSeq 3’UTR annotations (obtained from UCSC) and are associated with the corresponding genes. The rest of the unmapped ERs are then annotated to a) the corresponding gene if it is intronic, or b) the nearest gene upstream if it is within 5 kb from the gene ends. The intronic ERs are extended for 5 bp in each direction and the ERs downstream of genes are used to extend the gene’s 3’UTR annotation. The novel intronic polyA sites and extended gene annotations are then added to the gtf file used for gene counting. In total, additional internal polyA sites were added to 228 genes, and extended 3’UTRs were added to 476 genes, and 28 genes have both. Given the complexity of transcripts, the assignment to intronic polyA sites or 3’UTR extension may be impossible to distinguish in some cases. In addition, 2 of the 676 novel ERs did not make a difference to gene quantification as they overlapped with annotations from other genes.

### Gene quantification

Trimmed and filtered reads were assigned to genes using featureCounts (v1.6.2) (Liao et al., 2014) with parameters “ -s 1 -Q 255” for 3’UTR-seq.

### Processing of small RNA-sequencing reads

FastQC was used to examine the quality of sequenced reads. BBDuk (BBMap suite v37.90) was used to trim adapter sequences from reads with reference adapter sequences provided by BBMap suite and settings “hdist=1 mink=11” for small RNA-seq reads (Bushnell, 2014). For miRNA size specificity, only reads less than 23 nucleotides in length were retained. Following trimming, FastQC was used to examine the quality of trimmed sequenced reads. miRDeep2 mapper.pl was used with default parameters to map reads of at least 18 nucleotides in length to the mouse genome (mm10) (Friedländer et al., 2012). Known and novel miRNAs were identified using miRDeep2 main algorithm (miRDeep2.pl) with default parameters. For known miRNAs, the mature miRNA sequences in mouse were obtained from miRBase (v21) (Kozomara and Griffiths-Jones, 2014). For novel miRNAs, only those with miRDeep score ≥ 2 and a sequence not matching previously reported small RNAs (rfam alert = FALSE) were retained for downstream analysis.

### mRNA and miRNA normalization and differential analysis

All of the following analyses were performed in R (v3.6) unless otherwise specified. Low-count mRNAs and miRNAs were filtered out prior to analysis. Only mRNAs and miRNAs with normalized read count (counts per million mapped reads, CPM) > 2 in at least 10 samples were retained for downstream analysis. CPM was used because of the consistent read mapping with UTR-seq.

For mRNAs, quantification of mitochondrial genes was not considered in this study. Remove Unwanted Variation from RNA-Seq Data (RUV-seq, v1.18.0) was used with RUVg() function with empirically detected negative genes to estimate unwanted variations in mRNA data based on the previously shown superior performance of this method compared to methods using ERCC for library normalization (Risso et al., 2014). Empirical negative-control genes were identified with an ANOVA-like test comparing all conditions (FDR < 0.1). For miRNAs, RUV-seq was used with replicates (RUVs) to normalize and remove variation between batches from miRNA counts (Risso et al., 2014).

Quasi-likelihood F-test method was used to test for differential expression of mRNAs and miRNAs with a significance cutoff of absolute fold change (FC) > 1.5 and false discovery rate corrected (FDR) < 0.05 using edgeR (v3.26.5) (McCarthy et al., 2012; Robinson et al., 2010).

### Novel miRNA identification

The mature sequences of novel miRNAs which were included in the miRNA differential expression analysis were used for miRNA identification with “Single sequence search” function on miRBase (Kozomara et al., 2019) (https://www.mirbase.org/) with the following parameters: “Search sequences: Mature miRNAs”, “Search method: “BLASTN”, “E-value cutoff: 10”, “Maximum no. of hits: 100”, “Show results only from specific organisms: Mouse”, “Word size: 4”, “Match score: +5”, “Mismatch penalty: −4”. For miRNAs with more than one result, the miRNA with the best alignment (lowest E-value) is reported. If no miRNAs were identified in mouse, BLASTN was rerun with no species filter and the best alignment was reported. miRNAs with no results reported from any species are denoted with “NA”.

### Predicting transcriptional regulation using Lisa

Transcriptional regulator (TR) binding sites were predicted using Lisa (Qin et al., 2020) (http://lisa.cistrome.org/) with genes which were female- or male-biased in 2 or more ages between PD27, PD32, PD37. The full Lisa model was applied (“TR ChIP-seq Peak-RP (regulatory potential)” and “ISD-RP (in silico deletion-regulatory potential) for both motif and ChIP-seq” methods) using the DNase-seq and H3K27Ac ChIP-seq data and 3000 genes which were randomly selected as the background gene set. Results combined from H3K27ac-ChIP-seq and DNase-seq ISD models, and TR ChIP-seq peak-only models using the Cauchy combination test are shown for the ChIP-seq model.

### Pathway enrichment analysis

All pathway enrichment analyses were performed using g:ProfileR (v0.6.7) in R (v3.6). For differential expressed pathway enrichment, all detected genes in this dataset were used as background. For miRNA-gene target pathway enrichment, all gene targets detected in this dataset were used as background with parameters “min_set_size = 3, min_isect_size = 2”.

### Co-expression module identification

Gene co-expression modules were identified using CEMitool (Russo et al., 2018) (version 1.8.2) using log2-transformed, normalized read counts with default settings. Briefly, CEMitool first uses an unsupervised method to filter for genes with sufficient variation. By default, CEMitool models the variants of the genes as an inverse gamma distribution and chooses genes with a p value < 0.1. Next, it automatically determines the similarity criteria before it separates genes into modules using the dynamic tree cut method. Hub genes are identified by ranking the summed similarities between a certain gene and all other genes in the same module.

### miRNA-gene target correlation

Computationally predicted gene targets were curated from TargetScanMouse (v7.2) (Agarwal et al., 2015) and experimentally validated gene targets were curated from miRTarBase (v8.0) (Chou et al., 2018). Only miRNA-gene target pairs from TargetScan with “Cumulative weight context score” < −0.1 were used. “Context score” for a specific target site is defined by Agarwal et al. 2015 as the summed contribution from 14 features which likely influence miRNA targeting a given gene, including “site type”, “local AU”, “3’ UTR length”, and “Probability of conserved targeting” (full feature list http://www.targetscan.org/vert_70/docs/context_score_totals.html). Spearman’s correlation coefficient (rho) was calculated for each pair using log2-transformed normalized counts and p-values were adjusted with FDR to account for multiple testing. Pairs were considered negatively correlated if rho < 0 and FDR-adjusted P-value < 0.1.

### Integration of single-nuclei RNA-seq dataset from Ruf-Zamojski et al. 2021

Single-nuclei RNA-seq (snRNA-seq) of 10-12 week-old snap-frozen male and female C57BL/6 mouse pituitary gland was obtained from <u>GSE151961</u> and processed using Seurat 4.1.0 (Hao et al., 2021). Data was merged from replicates for each sex separately. Mitochondrial and ribosomal genes were filtered out of merged data. Filtered merged data for each sex was then normalized using SCTransform (Hafemeister and Satija, 2019). Data integration between sexes was performed using Seurat with the normalized merged data (Stuart et al., 2019). Cell types were labeled by comparing gene markers identified in cell clusters calculated by Seurat using the “FindAllMarkers” functions to gene markers identified in the original paper (Ruf-Zamojski et al., 2021). For all downstream analyses, the “Debris” cluster, which was also identified in the original paper, was removed. This analysis was performed using R version 4.1.2. For detailed methods, our code is available at https://github.com/wilsonlabgroup/pituitary_transcriptome_analyses.

### Cell-type enrichment of co-expression module genes

A one-sided Kolmogorov–Smirnov (KS) test was performed to compare the distribution of expression of a given co-expression module gene within each cell type versus its distribution of expression values in all other cell types based on the expression data from the Ruf-Zamojski et al. 2021 snRNA-seq dataset (see **Methods** for data processing details). Resulting *P*-values were FDR-adjusted for multiple testing. Only co-expression module genes with FDR-adjusted *P*-value ≤ 0.05 in at least one cell type comparison were plotted. A one-sided hypergeometric test was then used to determine if there was enrichment for a group of module genes with statistically greater expression based on the KS test (FDR-adjusted *P*-value ≤ 0.05) in each cell type. Resulting *P-*values were FDR-adjusted for multiple testing and a group of module genes was considered to be significantly enriched if FDR-adjusted P-value < 0.05. This analysis was performed using R version 4.1.2.

### Proportions in Admixture RNA-seq deconvolution

Proportions in Admixture RNA-seq deconvolution was run as part of a wrapper script in scMappR (v1.0.7) (Sokolowski et al., 2021) using the “compare_deconvolution_methods” function which calls the ADAPTS package (v1.0.21) (Danziger et al., 2019) with RUV-seq normalized bulk counts and a custom signature matrix. In scMappR, the Proportions in Admixture method is called “WGCNA”. The custom signature matrix was generated using the “generes_to_heatmap” function from scMappR with gene markers identified from sex-integrated Ruf-Zamojski snRNA-seq data (see **Integration of single-nuclei RNA-seq dataset from Ruf-Zamojski et al. 2021** for data processing) and selecting only the top 3000 genes with the greatest variance.

### Identifying cell-type-specific sex-biased genes using scMappR

Cell-weighted fold-change (cwFC) of PD37 sex-biased genes was calculated using scMappR (v1.0.7) (https://cran.r-project.org/package=scMappR) with sex-integrated adult C57BL/6 mouse pituitary transcriptome (Ruf-Zamojski et al., 2021) using the “WGCNA” (Proportions in Admixture) deconvolution method. Genes which were not detected in the single cell reference dataset were filtered out or were flagged as a “false positive” by scMappR (cwFoldchange_gene_flagged_FP) were filtered out. Genes were further filtered to include outliers as determined by scMappR based on their cwFC (cwFoldchange_gene_assigned). Finally, only genes with an absolute gene-normalized cwFC > 0.5 for a given cell-type were considered cell-type-specific sex-biased genes and shown in the heatmaps This analysis was performed using R version 4.1.2.

## Results

### Profiling postnatal mouse pituitary gland development with 3’UTR-seq and small RNA-seq

To assess changes in the mouse pituitary transcriptome across postnatal development, we profiled the pituitary RNA expression at five postnatal days spanning the pubertal transition (PD: 12, 22, 27, 32 and 37). We observed physical markers of pubertal onset, preputial separation (PS) and vaginal opening (VO) occurring on average at PD27 and PD29 in male and female mice respectively in our C57BL/6J colony ((Corre et al., 2016); Figure 1A). In all analyses performed (see Figure 1B for a summary of analysis workflow), pubertal onset refers to ages at which PS and VO were recorded.

**Figure 1.**
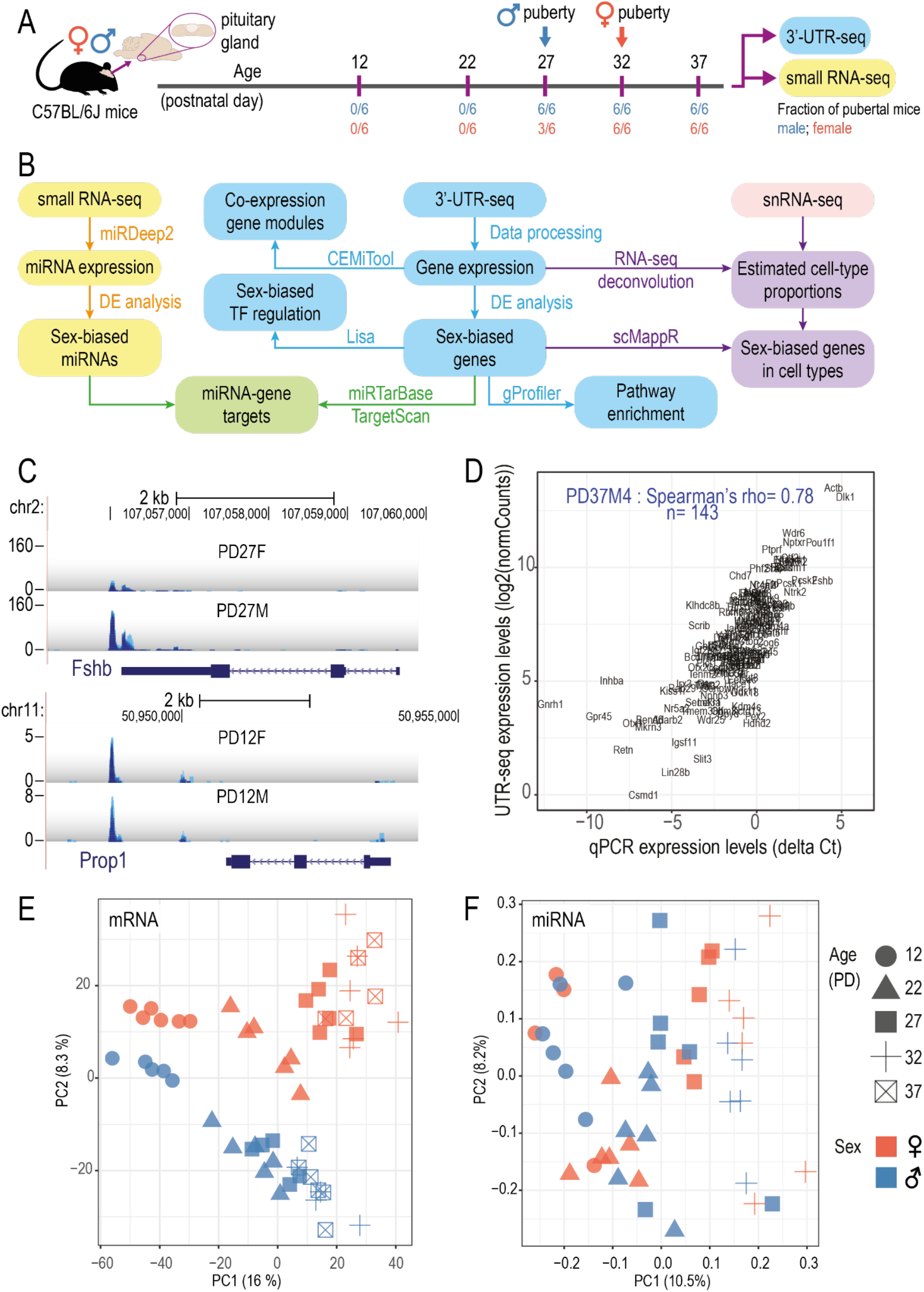
Overview of the pituitary transcriptome during postnatal development in male and female samples. **A.** Schematic of experimental design. Marks (purple) on the timeline denote the age in which the pituitary gland was collected. Vertical arrows denote average ages for onset of puberty of the specified sex in our colony (determined by preputial separation for males or vaginal opening for females). Fraction of pubertal mice out of total mice for males (blue) and females (red) at each age is shown. **B.** Schematic of analysis workflow. Summary of miRNA expression analyses (yellow), gene expression analyses (blue), miRNA-gene target identification (green), processing of single-nuclei RNA-seq (snRNA-seq) data from Ruf-Zamojski et al. 2021 (red), and combining snRNA-seq data with bulk gene expression data (purple). **C**. Genome browser screenshots showing QuantSeq signal at *Fshb* and *Prop1*. X-axis: genomic coordinates; y-axis: reads per million mapped reads (RPM); PD: postnatal day. Gene name and gene model are shown on the bottom of each panel. Each track represents overlapping signal from 5-6 biological replicates. **D.** Scatter plot showing the correlation between gene quantification measured by qPCR and by 3’UTR-seq in one pituitary sample (PD37M4). X-axis: ΔCt values obtained by qPCR; y-axis: log2-transformed normalized counts (log2(normCounts)) values obtained by 3’UTR-seq. Sample name, Spearman correlation coefficient, and number of genes included are labelled on the plot. PCA plot for pituitary gland samples based on **(E)** gene expression and **(F)** miRNA expression. Principal component analysis (PCA) was performed using log2(normCounts) after filtering for low-count genes/miRNAs and normalization using RUVseq. Only scores of the first 2 PCs are shown. Age is indicated by shape while sex is indicated by colours.

To measure mRNA expression in a genome-wide, cost-effective, and relatively high-throughput manner, we first automated the QuantSeq 3’ mRNA-Seq protocol which profiles 3’UTR of mRNA transcripts (**Methods** and **Supplementary figure S1A**). We then profiled between 5 and 6 biological replicates for each sex at each of the five postnatal days (55 libraries total, see **Supplementary table S1** for quality control metrics).

It was shown previously that 3’UTR profiling could miss the expression of a gene if the gene annotation does not capture novel or tissue-specific 3’UTRs. For example, *Prop1* expresses a novel 3’UTR in the pituitary and was missed in single-cell RNA-seq data generated using 10X Genomics’ Chromium technology (Cheung et al., 2018). To address this, we refined gene 3’-end annotations by identifying clusters of sequencing reads from our data and re-annotating 3’UTRs (**Methods** and **Supplementary figure S1A**). Improved pituitary-specific 3’UTR annotations were generated for 676 genes, allowing for assignment of significantly more reads to them, including important pituitary genes such as *Pou1f1*, *Ghrhr*, *Fshb*, and *Prop1* (Figure 1C**, Supplementary figure S1B**). UCSC genome browser tracks for the 3’UTR-seq signal of all samples are available at: https://genome.ucsc.edu/s/huayun/mouse_pituitary_utrseq.

To profile miRNA expression, we performed sRNA-seq in the same male and female samples used for 3’UTR-seq (PD: 12, 22, 27, and 32; n=5-6 biological replicates; 48 libraries total). Using the miRDeep2 workflow, we identified 273 known mouse miRNAs (miRBase v21) and 19 novel miRNAs (**Supplementary table S2**). The mature sequences of the novel miRNAs were used to identify homologous miRNAs in other species reported in miRBase v21 and precursor genome coordinates of the novel miRNAs were used to gain insight into their mechanism of biogenesis (Kim, 2005; O’Brien et al., 2018; Ruby et al., 2007) (see **Methods**).

For both 3’UTR-seq and sRNA-seq experiments, the biological replicates were well correlated (Pearson’s correlation coefficient: 3’UTR-seq 0.95-0.97; sRNA-seq 0.86-0.90) (**Supplementary figure S2A-B**). Furthermore, gene expression level quantified by 3’UTR-seq correlates well with microfluidic qPCR data previously generated from the same 55 RNA samples (178 puberty-related genes plus 5 control genes) (Hou et al., 2017) (median Spearman correlation coefficient: 0.74) (Figure 1D, **Supplementary figure S1C**). Using principal component analyses (PCA), we observed a separation of PD12 samples from PD22 and older for gene expression profiles along PC1 (Figure 1E). Although more subtle, we observed samples distributed by age along PC1 for miRNA expression profiles (Figure 1F). We also observed separation between male and female samples along PC2 at all ages based on gene expression profiles, which became more pronounced across postnatal age (Figure 1E). In contrast, no obvious sex differences were observed in our miRNA expression data (Figure 1F).

### Sex-biases in the transcriptome occur prior to puberty

We quantified sex differences in the pituitary transcriptome by comparing the expression of genes and miRNAs between male and female samples at each age. Across all the profiled ages, we observed an increase in the numbers of sex-biased genes and miRNAs, with the most dramatic increase occurring at PD27, when all the males and half of the females had gone through puberty (Figure 1A, Figure 2A-B; see Table 1, Table 2 for list of significant sex-biased genes and miRNAs; see **Supplementary table S3-4** for full list of differential analysis results).

**Figure 2.**
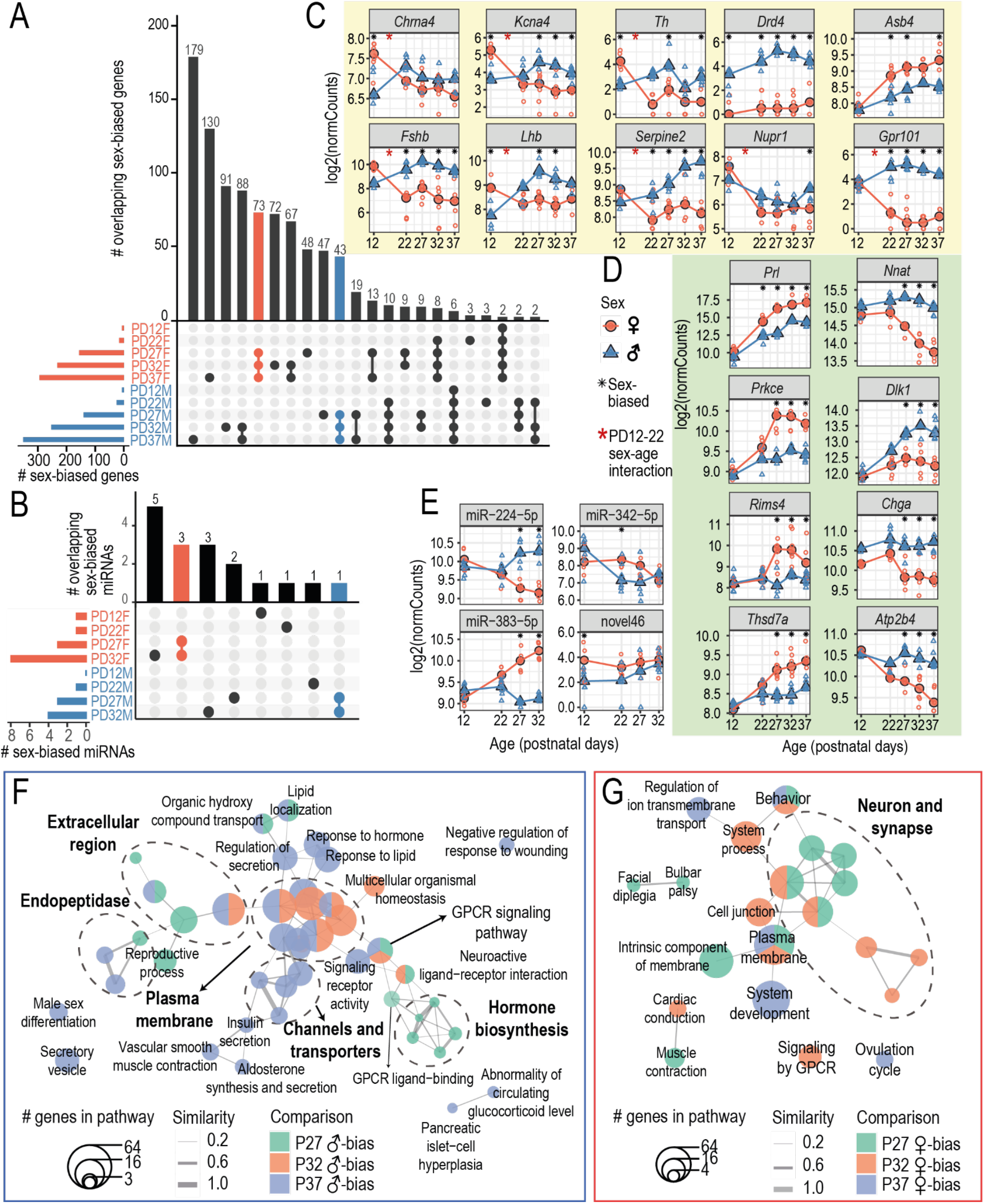
Pituitary transcriptome is increasingly sex-biased across postnatal development. Barplot showing the number of **(A)** intersecting sex-biased genes or **(B)** intersecting sex-biased miRNAs between each age. Horizontal bars on the bottom left side of each plot show the numbers of male- (blue) or female-biased (red) genes/miRNAs at each age (absolute FC > 1.5; FDR < 0.05). Different intersection combinations between sex-biased genes/miRNAs identified at each age are represented by the dotplot. The number of genes/miRNAs which intersect in the indicated combination of sex comparisons is shown by the vertical barplots (# overlapping sex-biased genes/miRNAs). Specifically, the coloured vertical bars represent the number of genes that are consistently female-biased (red) or male-biased (blue) at PD27, PD32, and PD37. Expression plots of example **(C)** pre-pubertal (PD12-22) sex-biased genes and genes with sex-by-age effect between PD12 and PD22, **(D)** peri-/post-pubertal sex-biased genes, and **(E)** sex-biased miRNAs. Log2-transformed normalized counts (log2(normCounts)) are plotted for each gene/miRNA. Expression changes are shown across ages (x-axis). Large, filled points represent median expression at each age and unfilled points represent each biological replicate. Blue: male samples; red: female samples. Black asterisks highlight ages at which the corresponding genes/miRNAs are detected as sex-biased and red asterisks higlight genes with significant sex-by-age effect between PD12 and PD22. Network representation of pathways enriched for **(F)** male-biased and **(G)** female-biased differentially expressed genes. Each node represents a pathway and nodes are connected based on similarity in genes found enriching for the connected pathways (Jaccard distance). Node size represents the number of differentially expressed genes enriching for the given pathway and nodes are coloured based on the differential expression comparison in which the genes were identified. Pathways that share similar genes are circled (dashed lines) and labelled manually based on pathway functions. Specific pathways and their manual labels can be found in **Supplementary table S5**.

**Table 1.**
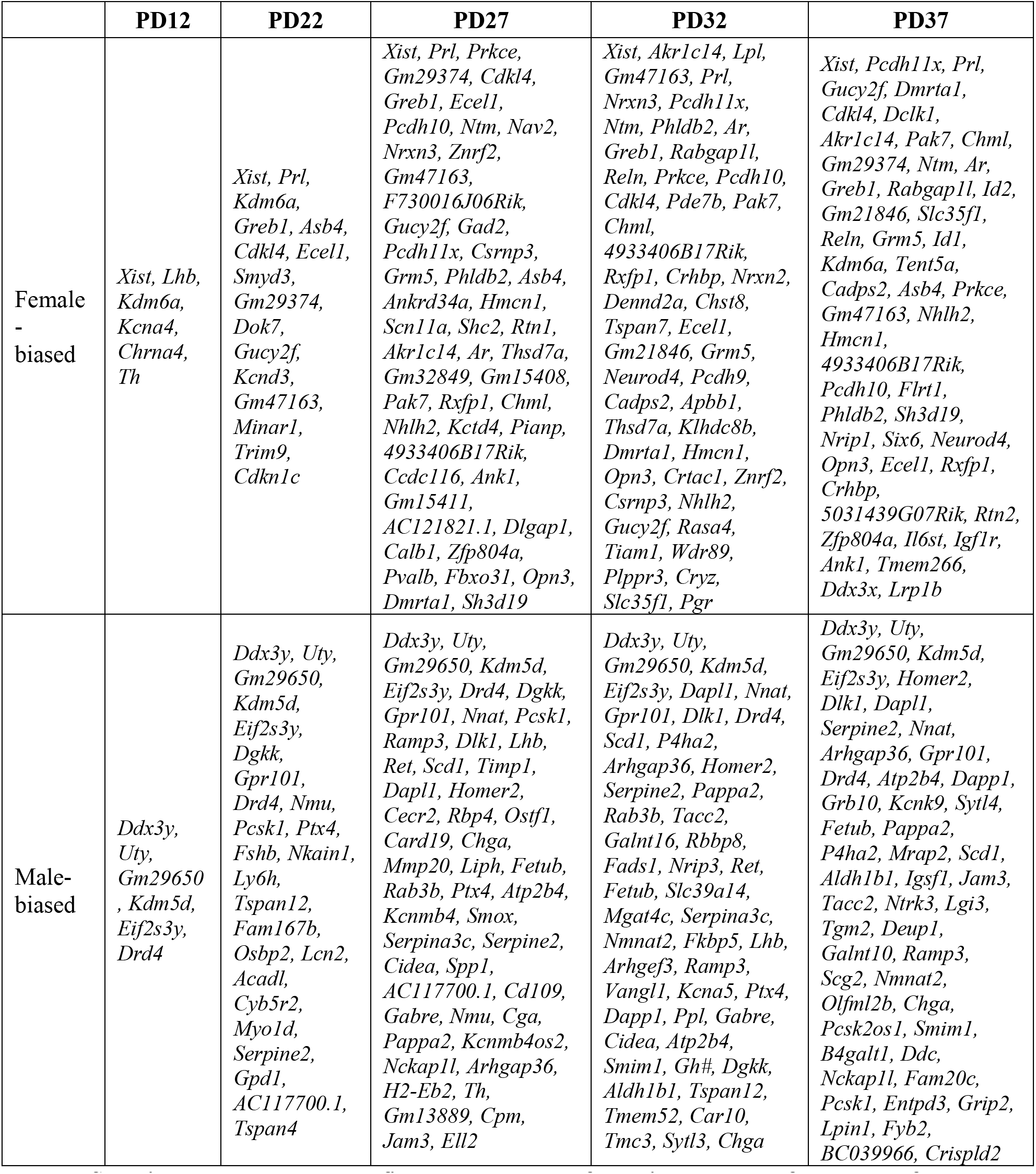
Sex-biased genes at each profiled age ranked by false discovery rate followed by fold-change (capped at 50 genes per comparison).

**Table 2.** Sex-biased miRNAs at each profiled age ranked by false discovery rate followed by fold-change. No results are shown for PD37 as this age was not profiled by sRNA-seq.

|  | PD12 | PD22 | PD27 | PD32 |
| --- | --- | --- | --- | --- |
| Female-biased | novel46 | miR-342-5p | miR-383-5p, miR-30d-5p, miR-17-5p | miR-383-5p, miR-205-5p, miR-146a-5p, miR-17-5p, miR-485-5p, miR-30d-5p, novel184, miR-669f-5p |
| Male-biased | NA | miR-499-5p | miR-224-5p, miR-181a-5p, miR-365-2-5p | miR-224-5p, novel37, miR-871-3p, miR-30b-5p |

Although we observed most sex-biased gene expression at peri-and post-pubertal ages, sex differences in the pituitary begin to manifest earlier in postnatal development at PD12 (Table 1). At PD12, 12 genes, including seven sex chromosome-linked genes: *Ddx3y*, *Uty*, *Kdm5d*, *Gm29650*, *Eif2s3y* (Y-linked), *Xist* and *Kdm6a* (X-linked); and five autosomal genes, *Chrna4*, *Kcna4*, *Lhb*, *Th*, and *Drd4*, are identified as significantly sex-biased (FDR < 0.05, absolute fold change > 1.5) (Figure 2C**, Supplementary figure S3A**). The seven sex chromosome-linked genes were consistently male-or female-biased across all the ages we profiled. Other than *Ddx3y*, *Eif2s3y*, *Uty*, *Xist*, and *Lhb* (Bjelobaba et al., 2015; Eckstrum et al., 2016), the other sex-biased genes we detected at PD12 have not been reported previously in pituitary.

We identified 25 male-biased and 16 female-biased genes at PD22 (Table 1), an age preceding puberty, including puberty-related genes *Dgkk*, *Fshb*, *Osbp2*, and *Pcsk1*, which we previously identified as being sex-biased using microfluidic qPCR (Hou et al., 2017) (Figure 2C**, Supplementary figure S3A**). Of these 41 sex-biased genes at PD22, we found 18 genes which start to exhibit sex-biased expression at PD22 and maintain the same sex-biased trend at all older ages we profiled, (i.e. *Asb4* and *Serpine2*) (Figure 2A, 2C, **Supplementary Figure S3A**), demonstrating that there is establishment of sex-biased expression in the pituitary prior to puberty.

We noticed some genes exhibited sex-biased changes between PD12 and PD22 (i.e. *Fshb*). We then specifically tested for sex-by-age interaction effect, and identified 13 genes (*Fshb*, *Pcsk1*, *Serpine2*, *Lhb*, *Chrna4*, *Nupr1*, *Dgkk*, *Steap3*, *Timp1*, *Knca4*, *Gpr101*, *2010007H06Rik*, and *Th*) with significant sex-by-age interaction effect between PD12 and PD22 (FDR < 0.05), all of which showed decreased expression in females and increased expression in males (**Supplementary figure S3A**). Several of these genes are associated with gonadotrope function: *Fshb* and *Lhb* are genes encoding for gonadotrope-secreted hormones, *Chrna4* is a marker for gonadotropes in rats (Fletcher et al., 2019), and *Nupr1* is known to be involved in embryonic gonadotrope development (Million Passe et al., 2008). *Gpr101* and *Serpine2* are have been previously shown to have sex-biased gene expression in gonadotropes isolated from juvenile mice but not adult mice (Qiao et al., 2016). However, the role of the remaining genes in the pituitary gland remains to be seen.

In contrast to the abundant mRNA expression, we identified 3 miRNAs with sex biased expression prior to puberty (PD12 or PD22). The one sex-biased miRNA at PD12 displayed female-biased expression and was one of 20 miRNAs (after normalization and filtering out low-count miRNAs; **Supplementary table S2**) not previously identified in mice based on miRBase v21 (we tentatively named it novel46; Figure 2E, Table 2**, Supplementary figure S3B**). We did not observe any alignments of novel46 to miRNAs in other species found in miRBase v21. Based on its miRNA precursor coordinates, novel46 is a mirtron (a miRNA which is spliced from a host gene intron) expressed from the last intron of calcium voltage-gated channel subunit alpha1 G (*Cacna1g*), located on chromosome 11 (**Supplementary table S2, Supplementary figure S3C**).

At PD22, we found that miR-499-5p displayed male-biased expression, and based on its miRNA precursor coordinates, miR-449-5p is a mirtron expressed from *Myh7b* (Table 2**, Supplementary figure S3B**). We also found that miR-342-5p displayed female-biased expression at PD22, and based on its miRNA precursor coordinates, miR-342-5p is a mitron expressed from *Evl* (Figure 2E, Table 2**, Supplementary figure S3B**). However, neither *Myh7b* nor *Evl* were detected as significantly sex-biased at any profiled age.

### Peri-and post-pubertal sex differences in gene expression reflect sex differences in pituitary endocrine functions

At peri-and post-pubertal stages we detected similar numbers of male-and female-biased genes. The number of sex-biased genes roughly doubled across the pubertal transition (140, 253, and 351 male-biased genes and 156, 232, and 294 female-biased genes at PD27, PD32, and PD37, respectively; Figure 2A, Table 1**, Supplementary table S3**). Many of these genes (43 male and 73 female-biased) showed sex biased gene expression throughout the pubertal transition (Figure 2A, 2D**, Supplementary figure S3A**). While we recovered genes with previously known sex-biased expression in pituitary, such as *Dlk1* and *Prl* (Bjelobaba et al., 2015; Cheung et al., 2013; Hou et al., 2017), many other genes with whose sex-biased expression has not been previously identified were found (Figure 2D, Table 1). For example, Neuronatin (*Nnat*), which is among the most abundant transcripts in the pituitary (Nishida et al., 2005b), showed strong male biased expression after PD22.

Male-biased genes are enriched for pathways related to hormone synthesis and secretion (Figure 2F**, Supplementary table S5**), including “*peptide hormone biosynthesis*” (PD27, p = 7.22e-06), “*regulation of secretion*” (PD37, p = 7.87e-03), and “*secretory vesicle*” (PD37, p = 4.9e-04); and pathways related to reproduction, including “*male sex differentiation*” (PD37, p = 4.76e-02) and “*reproductive process*” (PD27, p = 4.14e-02). In addition, male-biased genes are enriched for pathways associated with signaling receptors, ion channels, extracellular region, and plasma membrane, like “*G-protein coupled receptor signaling pathway*”, which is enriched at all three ages (PD27: p = 4.7e-02; PD32: p = 1.22e-03; PD37: p = 8.67e-03). These pathways highlight sex-biases in components important for cell signaling, particularly in pituitary endocrine cells which are known to be activated in a neuron-like manner, such as through ion channels and G-protein coupled receptors (Stojilkovic et al., 2017, 2010). Finally, pathways related to endopeptidase inhibitor activity are also enriched, including several serpine family genes (*Serpine2*, *Serpina3c*, and *Serpinb1a*). Whether these endopeptidase inhibitors are involved in sex-biased processing of peptide hormones or neuropeptides remains to be seen.

Female-biased genes, including *Stat5a*, *Cckbr*, *Slit2*, *Robo2*, *Nrip1*, *Nhlh2*, *Prl*, and *Pgr*, are enriched for “*ovulation cycle*” (PD37: p = 1.03e-02), linking female-biased pituitary genes to female-specific physiological processes. Notably, other female-biased pathways are predominantly neuron-related (Figure 2G**, Supplementary table S5**), which could be attributed to genes expressed in the posterior pituitary, which contains axons extended from the hypothalamus, or genes expressed in neuroendocrine cells (Robinson and Verbalis, 2011). Particularly, female-biased genes at all three ages, including *Crhbp*, *Prl*, *Calb1*, *Pgr*, *Pak7*, *Reln*, *Dmrta1*, *Prkce*, *Ar*, *Nhlh2*, *Grm5*, and *Cacna1c*, are enriched for “*behavior*” (PD27: p = 5.32e-03; PD32: p = 4.25e-02; PD37: p = 5.32e-03). Several of these genes, *Crhbp*, *Calb1* (J.-T. Li et al., 2017), *Prkce* (Hodge et al., 2002), *Cacna1c* (Moon et al., 2018), and *Grm5* (Shin et al., 2015) are related to the regulation of stress, which is sex-biased in its activity (Oyola and Handa, 2017). *Crhbp* encodes corticotropin-releasing hormone binding protein (CRH-BP), whose mRNA and protein levels are both higher in female mouse pituitary (Speert et al., 2002). CRH-BP inhibits ACTH secretion by binding to CRH and its expression is induced by stress resulting in a stronger attenuation of stress response in females (Stinnett et al., 2015). However, other than *Crhbp*, the sex-biased expression and the functions of these genes have not been studied in the pituitary.

For miRNAs, we detected 3 and 4 male-biased miRNAs and 3 and 8 female-biased miRNAs at PD27 and PD32 respectively (Figure 2B, Table 2**, Supplementary figure S3B, Supplementary table S4**). miR-224-5p, a mirtron of *Gabre*, and miR-383-5p, a mirtron of *Sgcz*, respectively displayed male- and female-biased expression levels that are consistently detected from PD27 to PD32 (Figure 2E, Table 2). miR-224-5p has been reported to promote ovarian granulosa cell (GC) proliferation with TGFβ1 stimulation (Yao et al., 2010). Meanwhile miR-383-5p was also shown to respond to TGFβ1 stimulation but inhibit GC proliferation (Wei and Gao, 2019; Yin et al., 2012). Given that TGFβ1 signaling is also a known inhibitor of lactotrope proliferation (Sarkar et al., 2005), it would be interesting to know if miR-224-5p and miR-383-5p have a role in TGFβ1-mediated lactotrope proliferation.

### Connecting sex-biased miRNAs to target genes with sex-biased expression

To evaluate post-transcriptional regulation of sex-biased changes in the pituitary gland by miRNAs, we identified computationally predicted or experimentally validated miRNA-target gene pairs that both exhibit sex-biased expression. Since miRNAs are usually predicted to repress target gene expression, we additionally required miRNA and target gene pairs to be significantly negatively correlated in expression across matched samples (Spearman’s rho < 0, FDR < 0.1) (see **Methods** for details) (Figure 3A**, Supplementary table S6**).

**Figure 3.**
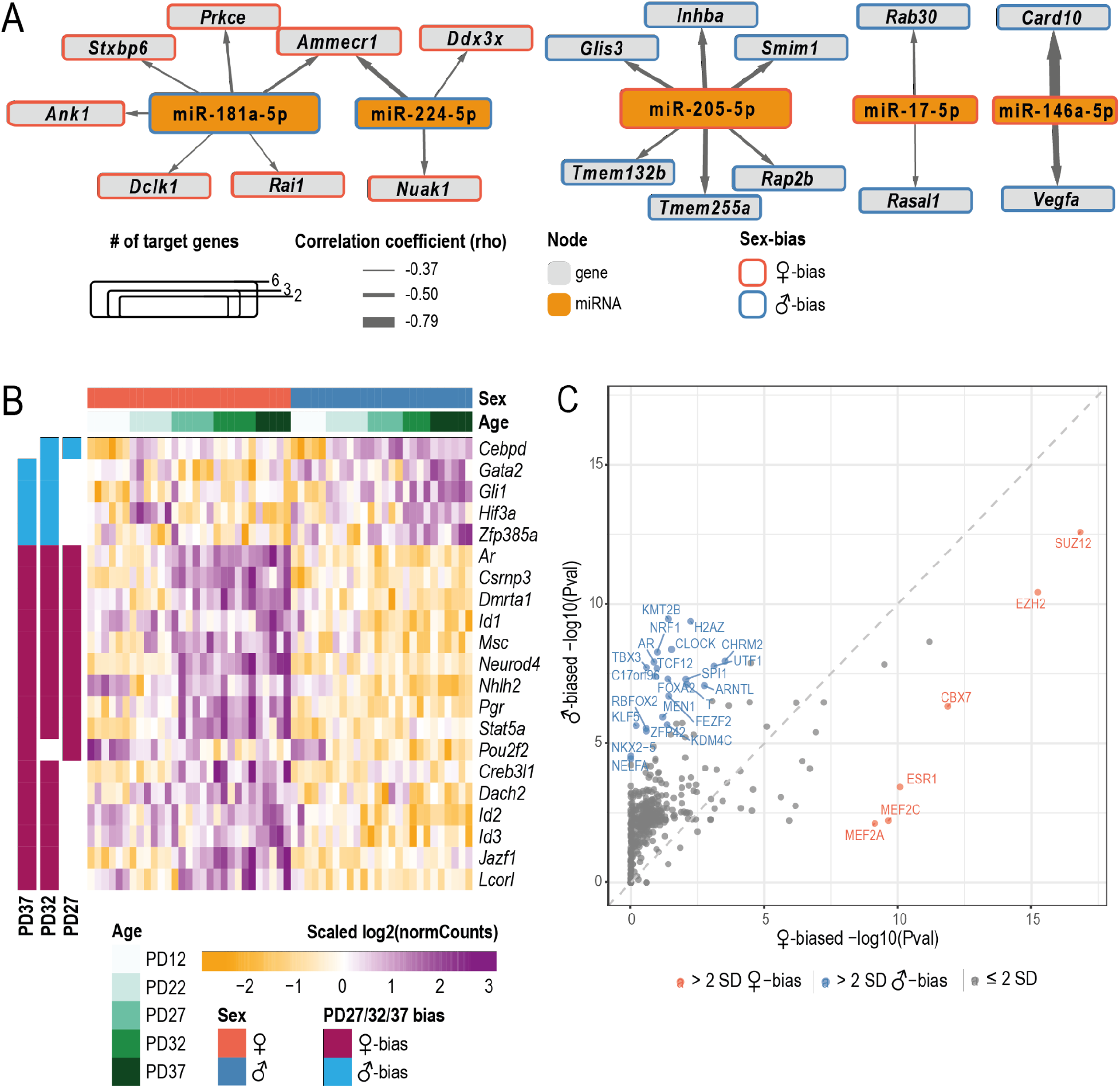
Regulation of sex-biased pituitary gene expression by miRNAs and transcriptional regulators (TRs). **A.** Interaction network of sex-biased miRNAs and their sex-biased target genes for all ages. Each node represents a gene (gray) or a miRNA (orange) with the node size representing its number of connections with other nodes. Outlines on nodes indicate gene/miRNA is female-biased (red) or male-biased (blue). Only miRNAs with 2 or more connections are shown. Edges between each node show a predicted interaction between the miRNA and gene. Edge thickness indicates Spearman’s correlation coefficient (rho) calculated for the given pair. **B.** Heatmap of sex-biased pituitary TR gene expression. TRs which are consistently sex-biased in at least two ages between PD27, PD32, and PD37 are plotted. Colors of the heatmap represent row-scaled and centered expression levels of each gene. Column annotation bars indicate sample age and sex. Row annotation bars indicate age at which the gene was found to be sex-biased. **C.** Scatterplot comparing Lisa TR rankings with combined P-values predicted to regulate female- and male-biased genes. Each TR is represented by a point. The combined −log10(*P*-value) is plotted for each TR based on female-biased and male-biased gene sets which are consistently sex-biased in at least two ages between PD27, PD32 and PD37. Coloured points show TRs which have change in −log10(*P*-value) between sexes which is two standard deviations greater than the mean change in −log10(*P*-value). Red points indicate TRs which are enriched for regulating female-biased genes; blue points indicate TRs which are enriched for regulating male-biased genes.

At pre-pubertal ages, PD12 and PD22, we did not identify any negatively correlated miRNA-gene pairs where the miRNA and its target gene were sex-biased. This is likely due to the relatively lower number of sex-biased genes and miRNAs identified at these pre-pubertal ages compared to post-pubertal ages.

At peri- and post-pubertal ages, we found that the male-biased miRNAs, miR-181a-5p (at PD27) and miR-224-5p (at PD27 and PD32), showed significant negative correlation with female-biased target genes, *Ammecr1* (target of miR-181a-5p and miR-224-5p), *Ank1* (target of miR-181a-5p), and *Prkce* (target of miR-181a-5p) (Figure 3A**, Supplementary table S6**). All three of these target genes have been shown to be estradiol-responsive (Dina et al., 2001; Kim et al., 2011) and are female-biased from PD27 onwards, suggesting a role for miR-181a-5p and miR-224-5p in regulating the pituitary response to increased circulating estradiol levels at puberty. Both *Ammecr1* and *Ank1* were previously identified as a target of miR-181a-5p in P13 mouse brain by Ago HITS-CLIP (Chi et al., 2009) and *Ammecr1* was previously identified as a target of miR-181a-5p in HeLA cells by microarray (Grimson et al., 2007). In comparison, female-biased miR-205-5p (at PD32) was negatively correlated with *Inhba* (Figure 3A**, Supplementary table S6**). *Inhba* encodes a subunit of inhibin, which is an inhibitor of FSH production in gonadotropes (Ling et al., 1986). Therefore, negative correlation of miR-205-5p to *Inhba* suggests that miR-205-5p may be relevant to sex biases in inhibin-regulated secretion of FSH from the pituitary gland.

### Predicting transcriptional regulators associated with sex-biased pituitary gene expression

Sex-biased expression of transcription regulators (TRs) is a principal mechanism by which sex-biased regulatory networks can be generated. Of the sex-biased genes we detected, 5 male-biased and 16 female-biased genes are annotated as TRs in mouse by AnimalTFDB3 (Hu et al., 2019) (Figure 3B). Known molecular functions in the pituitary and pituitary-related knockout phenotypes of these 21 sex-biased TRs are summarized in Table 3. Importantly, male-biased transcription factor gene *Cebpd* has been shown to inhibit PRL expression and lactotrope proliferation (Tong et al., 2011). In addition, *Gata2* and *Neurod4* are required for pituitary development (Charles et al., 2006; Zhu et al., 2006) and knockout of *Ar* or *Nhlh2* results in pituitary phenotypes including decreased preovulatory surge levels of LH and FSH (Wu et al., 2014) and hypoplastic pituitary (Cogliati et al., 2007).

**Table 3.**
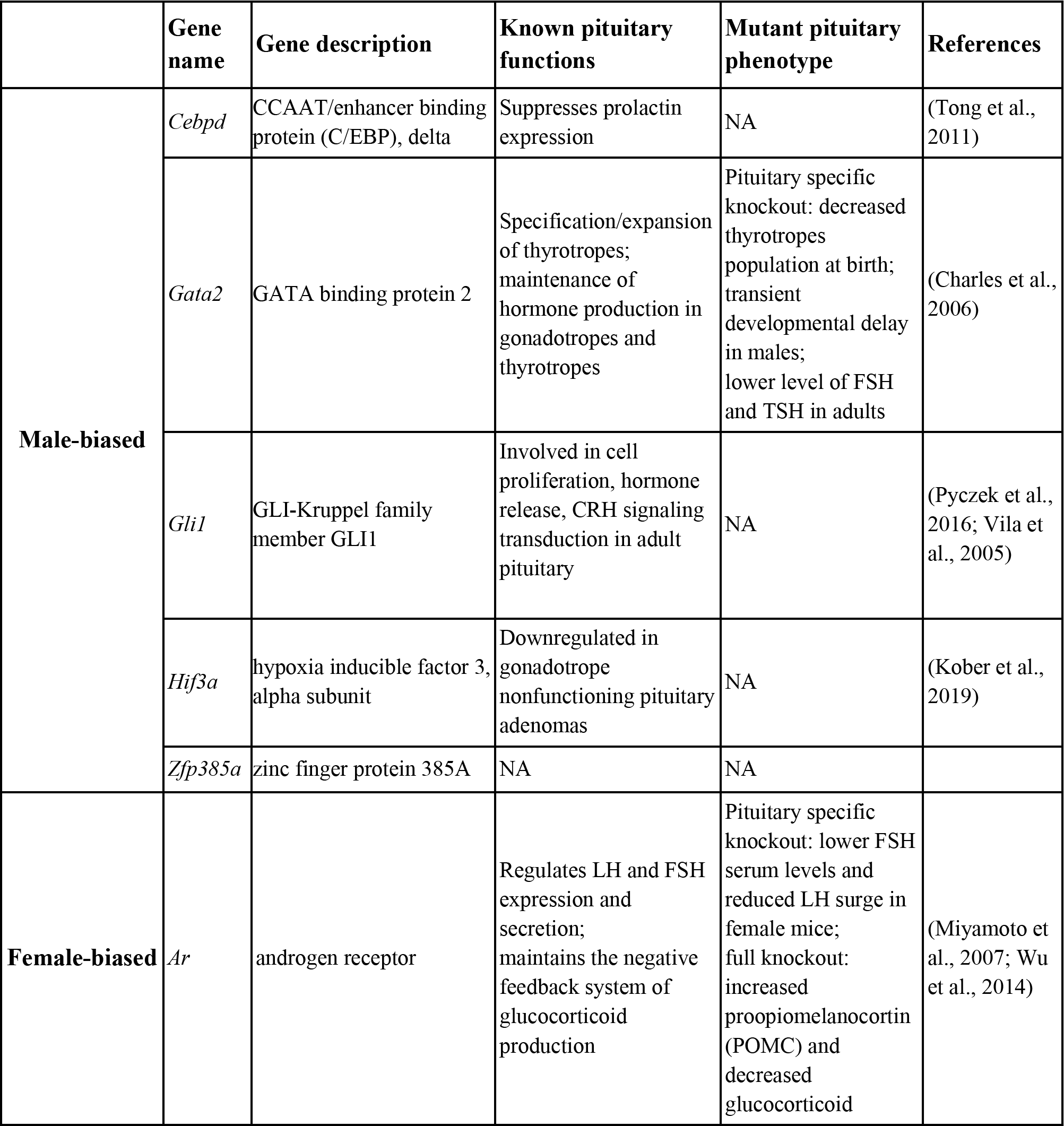
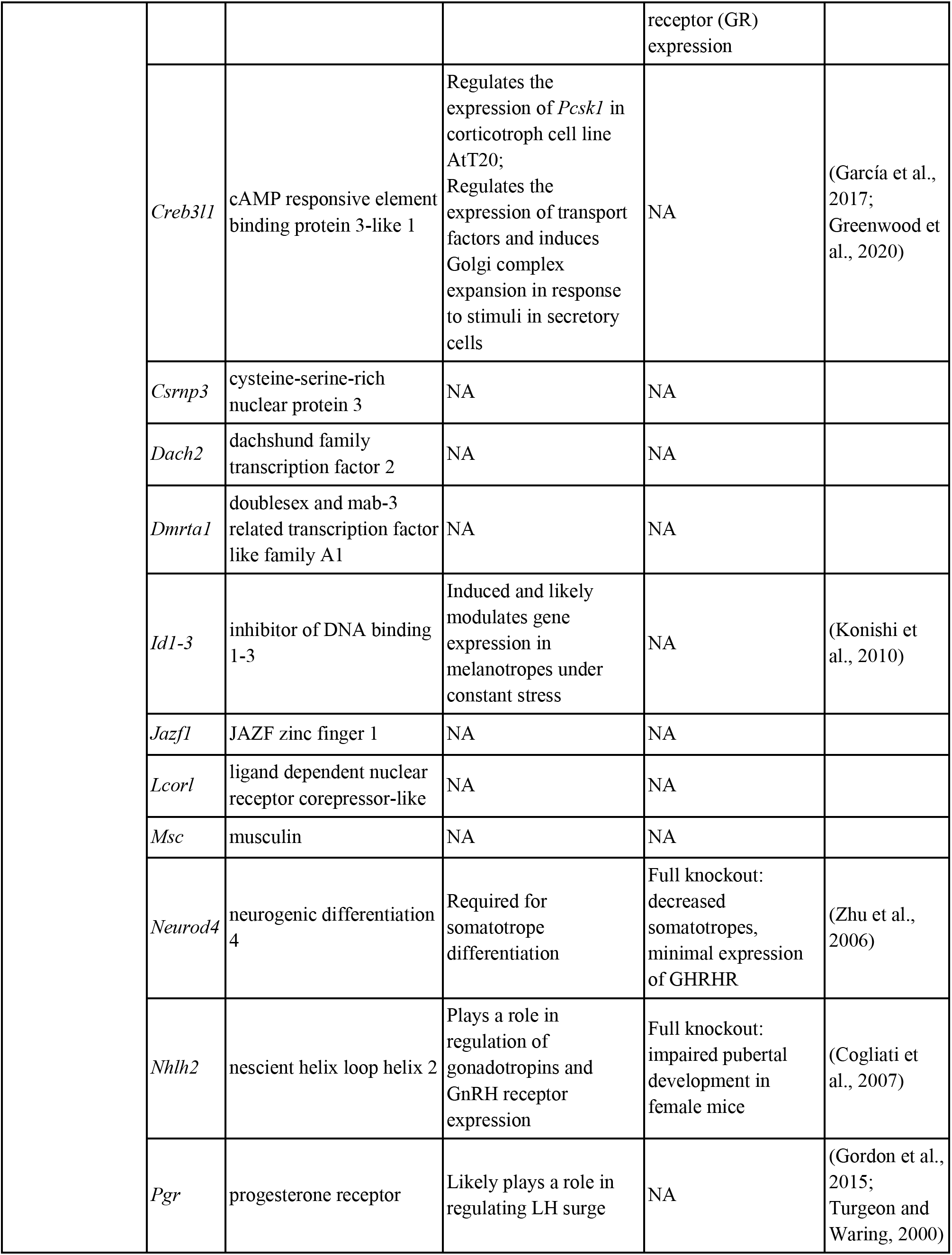
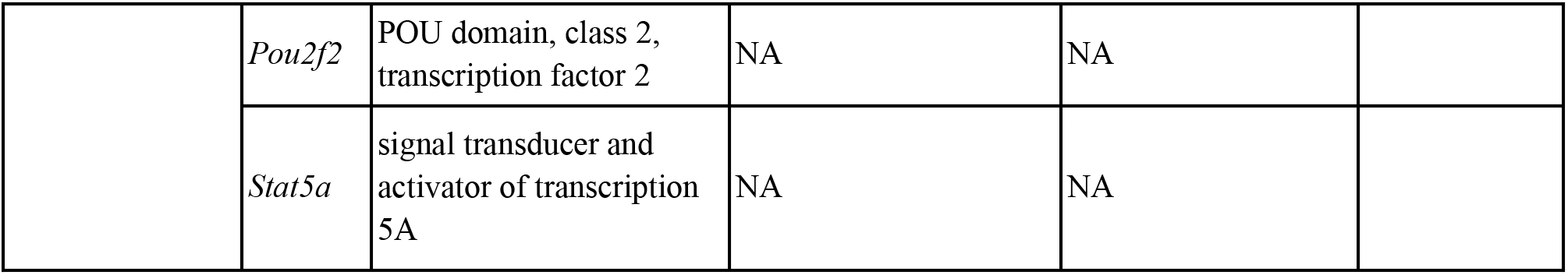
Pituitary-related phenotypes and functions of TRs that are sex-biased in at least two post-pubertal ages.

To gain insights into how tanscriptional regulatory proteins (TRs) might regulate sex-biased pituitary gene expression across pubertal transition, we used the bioinformatic tool Lisa (epigenetic Landscape In Silico deletion Analysis; (Qin et al., 2020)). Lisa takes a list of genes and builds transcriptional regulatory models based on both publicly available DNase and ChIP-seq datasets and returns predictions of candidate TRs that regulate them. We provide Lisa with pituitary genes classified as having sex-biased gene expression at two or more postnatal days between PD27, PD32, and PD37 in males (n=183) or females (n=174). We identified 6 and 22 TRs (Δ-log10(*P*-value) > mean ± 2SD) including gonadal nuclear hormone receptors, ESR1 and AR, that were predicted to regulate female-biased and male-biased genes respectively. In addition, CBX7, a component of Polycomb repressive complex 1 (PRC1), as well as SUZ12 and EZH2, components of Polycomb repressive complex 2 (PRC2), were predicted by Lisa to regulate female-biased genes. In contrast, a heterodimeric TR, CLOCK:BMAL1 (ARNTL), was predicted by Lisa to regulate male-biased gene expression (Figure 3C).

### Gene co-expression analysis reveals dynamic modules enriching for pituitary cell types

To further characterize developmental changes in the pituitary transcriptome, we utilized our temporal and sex-dependent gene expression profiling of the postnatal pituitary to build gene co-expression networks. Co-expressing genes tend to share related functions or regulatory pathways, and hence these networks are an effective approach to systemically explore mechanisms underlying sex-biased postnatal pituitary development. We used CEMiTool (Russo et al., 2018) to identify transcriptome-wide gene expression correlation networks in the postnatal pituitary transcriptome (see **Methods** for details). In total, nine co-expression gene modules were identified based on the expression of 1205 genes (Figure 4**, Supplementary figure S4-5**, **Supplementary table S7**).

**Figure 4.**
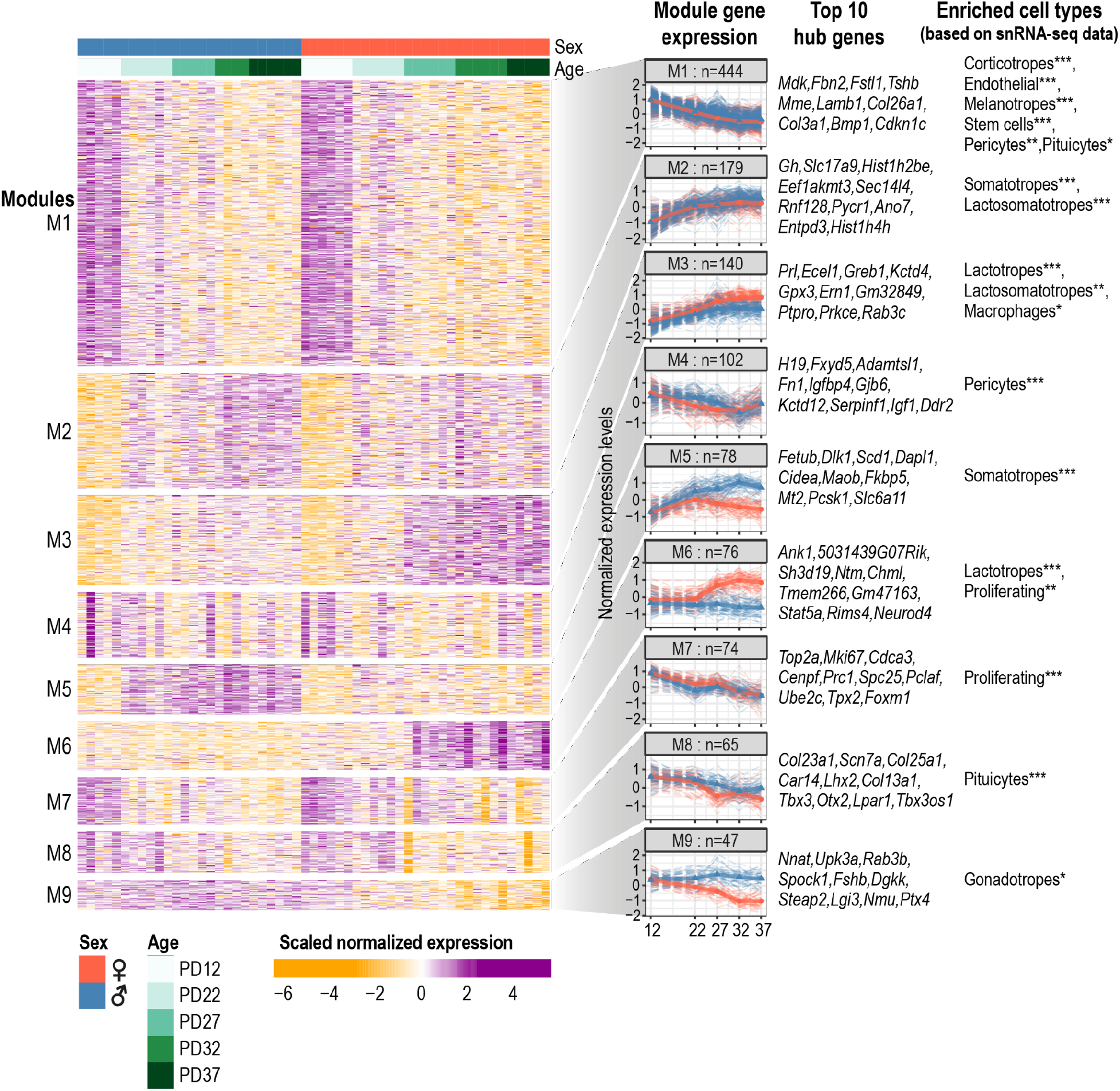
Co-expression network analysis identifies gene modules underlying pituitary transcriptome changes. Heatmap shows genes selected for co-expression analysis, separated into 9 modules. Column annotation bars indicate sample age and sex. Colours of the heatmap represent row-scaled and centered expression levels of each gene. A summary of the expression profiles of each module is shown in the left column of the right panel. The solid line represents the median expression (scaled and centered as shown in the heatmap) at each age and sex (red: female samples, blue: male samples) for all the genes in the corresponding module. Dash lines represent the scaled expression profiles of each gene in the module. Module names and number of genes included in the module are labelled. Top 10 hub genes (calculated based on genes’ connectivity within the modules) are listed for each module. All co-expressing module genes are listed in **Supplementary table S7**. Cell types in which module genes are enriched in based on single-nuclei RNA-seq expression from Ruf-Zamojski et al. 2021 are shown on the right (see **Supplementary figure S6C** and **Methods** for details). One-sided hypergeometric test: * FDR < 0.05, ** FDR < 0.01, *** FDR < 0.001.

The temporal expression profiles of the 9 co-expression gene modules showed three general patterns: 1) decreasing expression, particularly between PD12 and PD22 (Modules 1 (M1, M4, M7, and M8); 2) increasing expression (M2 and M3); and 3) sex-biased expression (M5, M6, and M9) (Figure 4**, Supplementary figure S5A-B**).

To further understand the nature of the co-expression modules, we used three independent approaches: 1) pathway enrichment analysis; 2) hub gene identification; and 3) an enrichment test where we ask if a set of genes in a module show enhanced expression in specific adult pituitary cell types (previously determined by single-nuclei RNA-seq (snRNA-seq) (Ruf-Zamojski et al., 2021)).

Using this approach, we found that our co-expression modules contain clear cell-type-enriched gene expression signatures. For example: 1) the M7 module, shows decreased expression during postnatal development in both males and females and is enriched for cell cycle-related pathways (**Supplementary figure S5C**); 2) The top M7 module hub genes include *Top2a*, *Mki67*, *Cdca3*, *Cenpf*, and *Prc1*, all of which are canonical markers of proliferating cells (**Supplementary figure S4**); and 3) Using published single-nuclei RNA-seq datasets of adult female and male mouse pituitary gland (Ruf-Zamojski et al., 2021) (**Supplementary figure S6A-B**), we found that there is a significant enrichment of M7 genes in the proliferating cell population (Hypergeometric test, FDR=2.60e-41) (Figure 4, **Supplementary figure S6C,** see **Methods** for details).

Similarly, the top hub genes for M2 and M5 include *Gh* and *Dlk1* (Figure 4), markers of somatotropes (Cheung et al., 2013), and there was a significant enrichment of both M2 (FDR =6.84e-11) and M5 (FDR=2.82e-10) module genes in somatotropes (**Supplementary figure S6C**). The top hub gene of M3 is *Prl*, a marker of lactotropes, and there is a significant enrichment of M3 module genes (FDR=3.18e-24) in the lactotropes. Finally, the top hub genes in M8 consist of pituicyte markers, including *Col13a1*, *Scn7*, and *Col25a1* (Chen et al., 2020), there is also a significant enrichment of M8 module genes (FDR=1.61e-17) within the pituicytes (**Supplementary figure S6C**). Overall, we observe a clear association between cell types and specific gene co-expression modules identified in the pituitary gland.

### Sex differences in cell proportions emerge prior to and across puberty

Although sex differences in the number of somatotropes, lactotropes and gonadotropes have been documented in the adult pituitary gland (Ho et al., 2020; Ruf-Zamojski et al., 2021; Sasaki and Iwama, 1988), much less is known about when these sex differences arise during postnatal development. To assess whether sex differences in cell proportions of somatotropes, lactotropes and gonadotropes are dynamic across pubertal development, we estimated cell-type proportions in our temporal bulk RNA-seq using the Proportions in Admixture RNA-seq deconvolution algorithm (Langfelder and Horvath, 2008) from Automated Deconvolution Augmentation of Profiles for Tissue Specific cells (ADAPTS) (Danziger et al., 2019) (Figure 5A, Figure 5B, **Supplementary figure S6D**). All cell types detected in the sex-integrated single-nuclei adult female and male pituitary transcriptome were used as our reference (Ruf-Zamojski et al., 2021) (**Supplementary figure S6A-B**).

**Figure 5.**
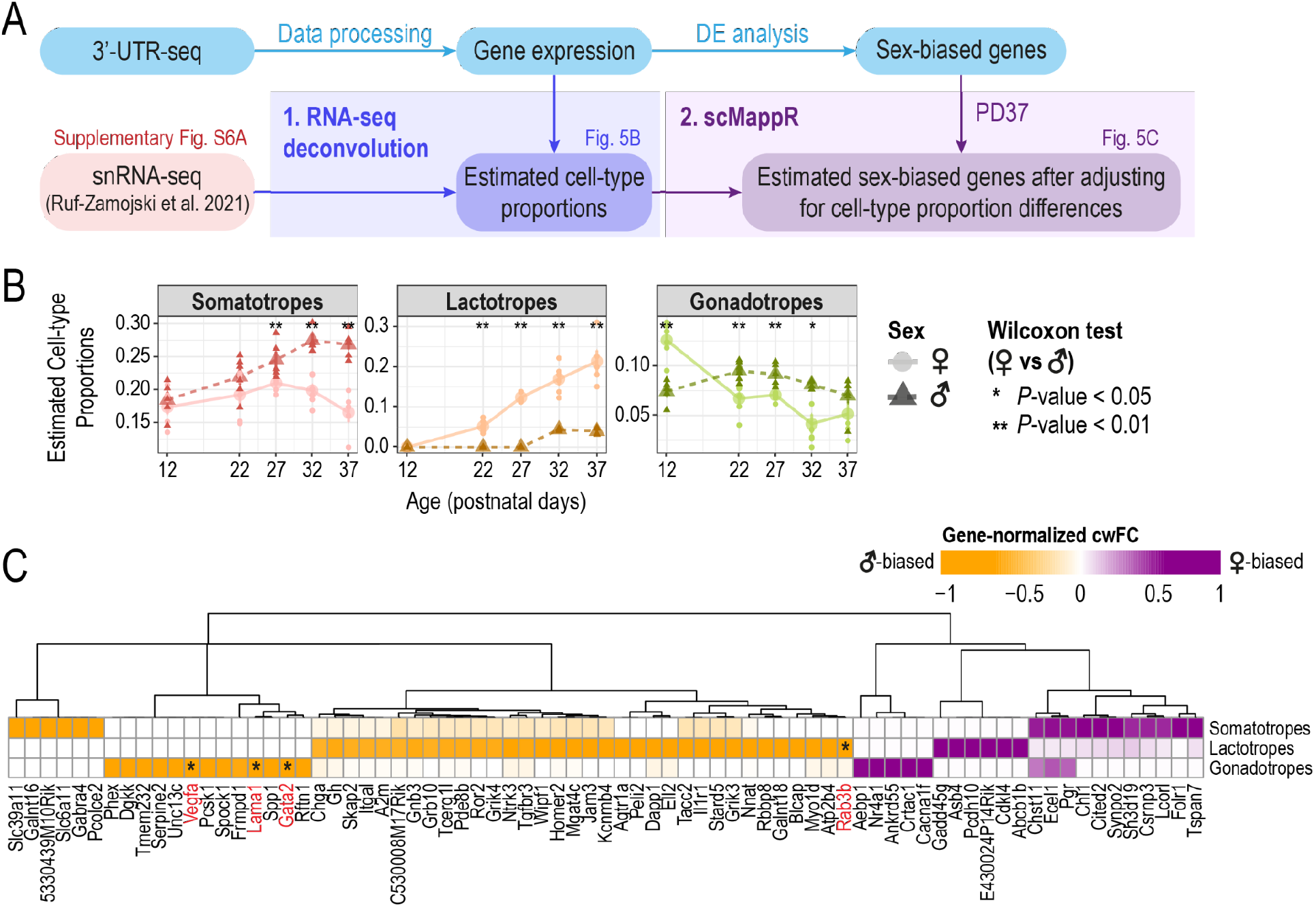
Estimating sex differences in pituitary cell types by leveraging single-nuclei RNA-seq. **A.** Schematic of analysis workflow for estimating sex differences in pituitary cell types using bulk gene expression with single-nuclei RNA-seq data from Ruf-Zamojski et al. 2021. **B.** Estimated cell-type proportions by RNA-seq deconvolution using Proportions in Admixture (WGCNA) changes across profiled ages of cell types with previously established sex-biased proportions in the adult pituitary. Estimated cell-type proportions are plotted across postnatal ages along the x-axis. Large circles and triangles represent the mean cell-type proportion at each age and small circles and triangles represent each biological replicate. Lighter colour, solid line, circle points: female samples; dark colour, dotted line, triangle points: male samples. Wilcoxon test was performed to compare cell proportions between both sexes at each age (\**P*<0.05, \*\**P*<0.01). See **Supplementary S6D** for all other pituitary cell type proportions. **C.** Heatmap of cell-weighted fold-changes (cwFC) for sex-biased genes at PD37 in somatotropes, lactotropes, and gondaotropes. Colour gradient indicates gene-normalized cwFC value calculated by scMappR; purple: more female-biased; orange: more male-biased. Genes with |gene-normalized cwFC| > 0.5 in at least one cell type are plotted. Genes shown to be sex-biased in the same direction in the same cell-type by (Fletcher et al., 2019) are indicated by the red text and an asterisk within the cell of the corresponding cell-type.

We found that we were able to recapitulate known sex biases in adult pituitary cell-type proportions (Ho et al., 2020; Ruf-Zamojski et al., 2021; Sasaki and Iwama, 1988) at our oldest profiled age, PD37, where the estimated cell-type proportions for somatotropes were significantly male-biased, and lactotropes were significantly female-biased (Figure 5B). In addition, we identified sex differences in estimated cell-type proportions which were dynamic across our earlier profiled ages. For example, we observed significant sex differences in estimated gonadotrope cell-type proportions between PD12 and PD22, ages prior to puberty, and these proportions show a sex-by-age trend (decreasing proportions from PD12 to PD22 in females and increasing in males), which could contribute to some of the sex-by-age bulk gene expression we observed (Figure 5B). At PD27, when puberty has occurred in a subset of our mice (Figure 1A), we observed the emergence of male-bias in somatotrope cell proportions and female-bias in lactotrope cell proportions (Figure 5B). The emergence of sex differences in relative cell proportions at ages leading up to and including puberty suggest that pubertal transition may be a key time window for the establishment of sex-biased proportions of hormone-producing cells observed in adult mice (Ho et al., 2020; Ruf-Zamojski et al., 2021; Sasaki and Iwama, 1988).

### Inferring sex-biased gene expression in pituitary cell types

We next examined if any sex-biased genes identified remained sex-biased after adjusting for cell-type proportion sex differences in somatotropes, lactotropes, and gonadotropes. To do this, we used a bioinformatic pipeline, Single-cell mapper (scMappR) (Sokolowski et al., 2021) to calculate a cell-weighted fold-change (cwFC) for each sex-biased gene identified in our bulk RNA-seq data (Figure 5A). Briefly, cwFCs were calculated for each sex-biased gene in somatotropes, lactotropes, and gonadotropes, by readjusting the bulk gene expression fold change with the cell-type specificity of the gene and the ratio of deconvolution-estimated cell-type proportions (determined by snRNA-seq). To minimize potential confounding effects from differences in ages, we focused on sex-biased genes identified at PD37, which is the closest age to the snRNA-seq reference dataset from adult male and female mice (Ruf-Zamojski et al., 2021). We identified 75 sex-biased genes which remain sex-biased after adjusting for sex differences in cell-type proportions (|cwFC| > 0.5) across somatotropes, lactotropes, and gonadotropes (Figure 5C, **Supplementary table S8**). Of these 75 sex-biased genes, 4 genes (*Gata2*, *Rab3b*, *Vegfa*, and *Lama1*) were found to be concordantly sex-biased in the corresponding cell type in the rat anterior pituitary (genes highlighted in Figure 5C, (Fletcher et al., 2019)).

Several of these genes have plausible links to regulating sex-biases in specific pituitary cell types. For example, the male-biased expression of *Rab3b* in lactotropes is consistent with its inhibitory role of PRL secretion in males (Perez et al., 1994). In addition, the male-biased expression of *Gata2* in gonadotropes is consistent with male-specific growth deficiency observed with pituitary-specific *Gata2* KO mice (Charles et al., 2006) and the male-specific reduction in serum FSH in gonadotrope-specific *Gata2* KO mice (Schang, 2020). Our analysis also reveals potential novel sex-biased roles for genes. For example, *Nr4a1* may be female-biased in gonadotropes, and this gene has previously been shown to be upregulated in response to GnRH (Ruf-Zamojski et al. 2018). Thus, by combining our temporal postnatal bulk pituitary 3’UTR-seq with a snRNA-seq dataset we can infer postnatal pituitary cell-type specificity of sex-biased genes and gain insights into sex-biased temporal gene regulation in a cell-type-specific manner.

## Discussion

The pituitary gland displays sex differences in its regulated physiological functions, including stress response, somatic growth, reproduction, and pubertal timing. By comparing pituitary gland gene and miRNA expression profiles between male and female mice during postnatal development, we identified sex-biased genes and miRNAs that contribute to sex differences postnatal pituitary development and function.

While the majority of sex differences in pituitary gene expression were observed at puberty and beyond, we observed robust sex-biased expression of genes and miRNAs prior to onset of physical markers of puberty (VO/PS) at PD12 and PD22. These sex differences may be attributed to irreversible organizational effects (McCarthy and Arnold, 2011) that are established by gonadal hormone exposures during fetal development and perhaps in the first week after birth in mice (R. Li et al., 2017). Unlike activational effects, sex differences established by organizational effects persist even when gonadal hormones are no longer present in the system, such as at pre-pubertal ages. We found that many sex-biased genes identified between PD12-PD22 are expressed in the gonadotropes, including *Fshb* and *Lhb*. We also predicted by RNA-seq deconvolution that gonadotrope cell proportions are male-biased prior to puberty. As gonadotropes produce LH and FSH to regulate gonadal maturation during puberty, these pre-pubertal sex differences in gonadotrope cell proportions and sex-biased expression suggest that these sex differences are necessary to prime the pituitary gland for its regulation of gonadal maturation during puberty.

By identifying a list of sex-biased genes we were able to predict trans regulators that are likely to underly their expression. Expectedly, we predicted that ESR1 and AR are involved in regulating female- and male-biased gene expression. We also predicted that female-biased gene expression involved regulation by polycomb complexes PRC1 (CBX7) and PRC2 (SUZ12 and EZH2). This observation is consistent with results obtained from the GTEx consortium who compared adult male and female pituitary gene expression (Oliva et al., 2020). Mechanistically, sex-biased deposition of H3K27me3 marks by PRC2 through its catalytic subunits Ezh1/Ezh2 has been shown to repress female-biased expression in the male adult mouse liver (Lau-Corona et al., 2020) and is also involved in the regulation of pubertal onset in female rats by inhibiting *Kiss1* expression in the arcuate nucleus to prevent the premature initiation of puberty (Lomniczi et al., 2013). Our study highlights that future studies of these mechanisms in the mouse pituitary would be ideal to study prior to puberty.

miRNAs have potentially important roles in responding to sex-biased hormonal release across multiple hypothalamic-pituitary axes. For example, multiple miRNAs have been shown to be estrogen-responsive in the neonatal mouse hypothalamus by the administration of an aromatase inhibitor (Morgan and Bale, 2017). miR-1948 and miR-802, identified as sex-biased in adult mouse liver, are known to be regulated by sex-biased pattern of growth hormone release from the pituitary gland (Hao and Waxman, 2018). Here we identified two male-biased miRNAs, miR-181-5p and miR-224-5p with estrogen-responsive predicted targets: protein kinase Cε (encoded by *Prkce*) signaling in response to epinephrine-induced inflammatory pain was found to be suppressed by estrogen in rats (Dina et al., 2001), and *Ammecr* and *Ank1* gene expression was upregulated by estradiol in mouse pituitary (Kim et al., 2011). We also identified a female-biased novel miRNA at PD12, novel46, whose host gene, *Cacna1g*, has been shown in anterior pituitary primary cells to promote LH secretion by estradiol signaling through estrogen receptor 1 (ESR1) upon GnRH-induction (Kim et al., 2011). While additional functional studies of these miRNAs is needed, these putative miRNA-mRNA connections represent potential gene regulatory networks underlying sex differences in postnatal pitutary gland development.

Sex differences in the number of somatotropes and lactotropes in the adult pituitary have been known for more than two decades (Sasaki and Iwama, 1988). These findings have recently been expanded on by single-cell genomic analyses of the adult pituitary (Fletcher et al., 2019; Ho et al., 2020; Ruf-Zamojski et al., 2021). In this study, we combined our temporal bulk RNA-seq data with single-cell RNA-seq data from adult pituitaries to to estimate sex differences in cell-type proportions during postnatal development. We suggest that these sex differnces in cell proporions occur prior to puberty. There are several proposed mechanisms by which cell composition changes arise in the pituitary gland: differentiation from the adult pituitary stem cell niche (Andoniadou et al., 2013), transdifferentiation from differentiated cell-types (Ho et al., 2020; Scheithauer et al., 1990), and self-proliferation from existing cell-types (Oishi et al., 1993; Taniguchi et al., 2002). Further work profiling female and male pituitaries using single-cell genomic technologies during early postnatal development will be valuable as will experiments designed to perturb and study the impact that sex biased gene regulatory networks have on the cell composition of the pituitary gland.

By using bulk mRNA and miRNA profiling of multiple replicates of male and female mice at multiple postnatal timepoints we identified sex biases in gene expression which are above and beyond what would be expected by differences in cell type proportion. Nevertheless, our observation of sex differences in gene expression will greatly benefit from single cell genomic assays that profile RNA expression and chromatin accessibility during postnatal development. Early evidence of the power of these approaches for studying gene regulatory networks active in postnatal pituitary comes from Ruf-Zamojski et al. who used snRNA-seq and snATAC-seq to characterize the sex-biased specific regulatory landscape of the male (n=3) and female (n=3) adult mouse pituitary (10-12 weeks) (Ruf-Zamojski et al., 2021). Particularly, Ruf-Zamojski et al. highlighted a latent variable (LV) showing increased expression/accessibility in females. We found that 10 of the top 30 genes associated with this LV also show evidence of female biased expression in our study (*Ankra2*, *Crhbp*, *Ddx21*, *Ern1*, *Gadd45g*, *Greb1*, *Npr2*, *Nrg4*, *Rps6ka2*, *Stat5a*). Similarly, technological advances in single-cell small RNA sequencing (reviewed in (Hücker et al., 2021)), and in particular single-cell miRNA-mRNA co-sequencing techniques (Wang et al., 2019), will likely permit a higher-throughput way to assign miRNAs to specific cell types and study relationships with their target genes.

We performed a comprehensive profiling of pituitary gene and miRNA expression across the pubertal transition, a major postnatal developmental milestone, in both male and female mice. We have made this resource feely available so that others can use this data to design experiments to further understand the biology of sex differences in the pitutiary gland. Our intial analysis of these data identified novel pituitary-expressed sex-biased genes and miRNAs that cannot be explained by differences in cell-type proportions. We also revealed dramatic developmental changes between PD12 and PD22, the mechanisms of which remain to be seen. By providing evidence for pre-pubertal sex differences in pituitary cell-type proportions, a specific challenge becomes dissecting the mechanisms which generate these differences. The unique role of the pituitary gland in reproductive and stress-related processes that differ between sexes highlights the importance of such future work.

## Ethics approval

All studies and procedures were approved by the Toronto Centre for Phenogenomics (TCP) Animal Care Committee (AUP 09-08-0097) in accordance with recommendations of the Canadian Council on Animal Care, the requirements under Animals for Research Act, RSO 1980, and the TCP Committee Policies and Guidelines.

## Availability of data and materials

The datasets generated and analysed during the current study are available in the ArrayExpress repository under accession numbers: E-MTAB-9459, E-MTAB-9460. All datasets and scripts used to perform analyses in the paper and reproduce figures can be found here: https://github.com/wilsonlabgroup/pituitary_transcriptome_analyses.

## Authors’ contributions

Experimental and analysis study design was conceived by HH, C.Chan, DS, MFM, AG, ZZ, MRP, and MDW. Bioinformatic analysis was performed by HH, C.Chan and DS. Experiments were performed by KEY, AR, LUR, MH, C.Corre. Pipeline for preprocessing 3’UTR-seq data was developed by RQ and HH. The manuscript was written by HH, C.Chan, and MDW with support from all authors. AG, ZZ, MRP and MDW supervised and obtained funding for this work. All authors have read and approved the manuscript for publication.

## Acknowledgements

We would like to thank: Theresa Ten Eyck (Agilent), Jekaterina Aleksejeva and Stephanie Bannister (Lexogen), Karen Ho and Sergio Pereira (The Centre for Applied Genomics (TCAG)) for their efforts and guidance setting up the automation of the QuantSeq protocol; and Sunyun Lee for assistance with setting up the Shiny app.

