## Supplementary Figures for "Postnatal developmental trajectory of sex-biased gene expression in the mouse pituitary gland"

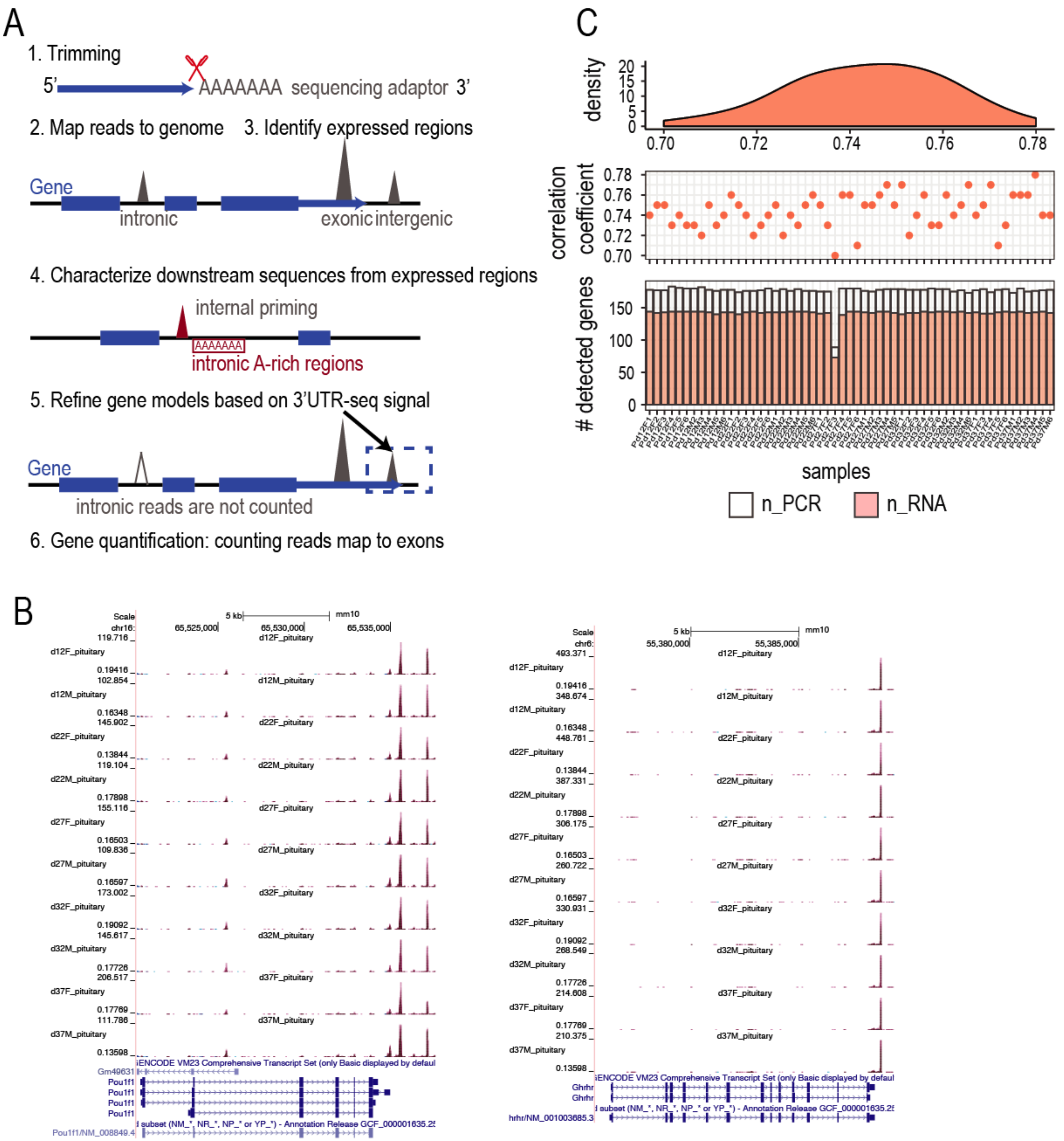


**Supplementary figure 1. Illustration of 3'UTR-seq method and quality control.**

**A.** Overview of the QuantSeq data analysis pipeline. **B.** Genome browser screenshot of extended 3'UTRs for genes *Pou1f1* and *Ghrhr*. Gene name and gene model are shown on the bottom of each panel. Each track represents overlapping signal from 5-6 biological replicates. **C**. Summary of qPCR vs. 3'UTR-seq comparisons across samples in all samples. Bottom panel: bar plots showing the numbers of genes detected using qPCR (white bars) and 3'UTR seq (red bars). The Spearman correlation coefficients between two experiments are shown for each sample in the dot plot (middle panel) and as well as in a density plot (top panel).

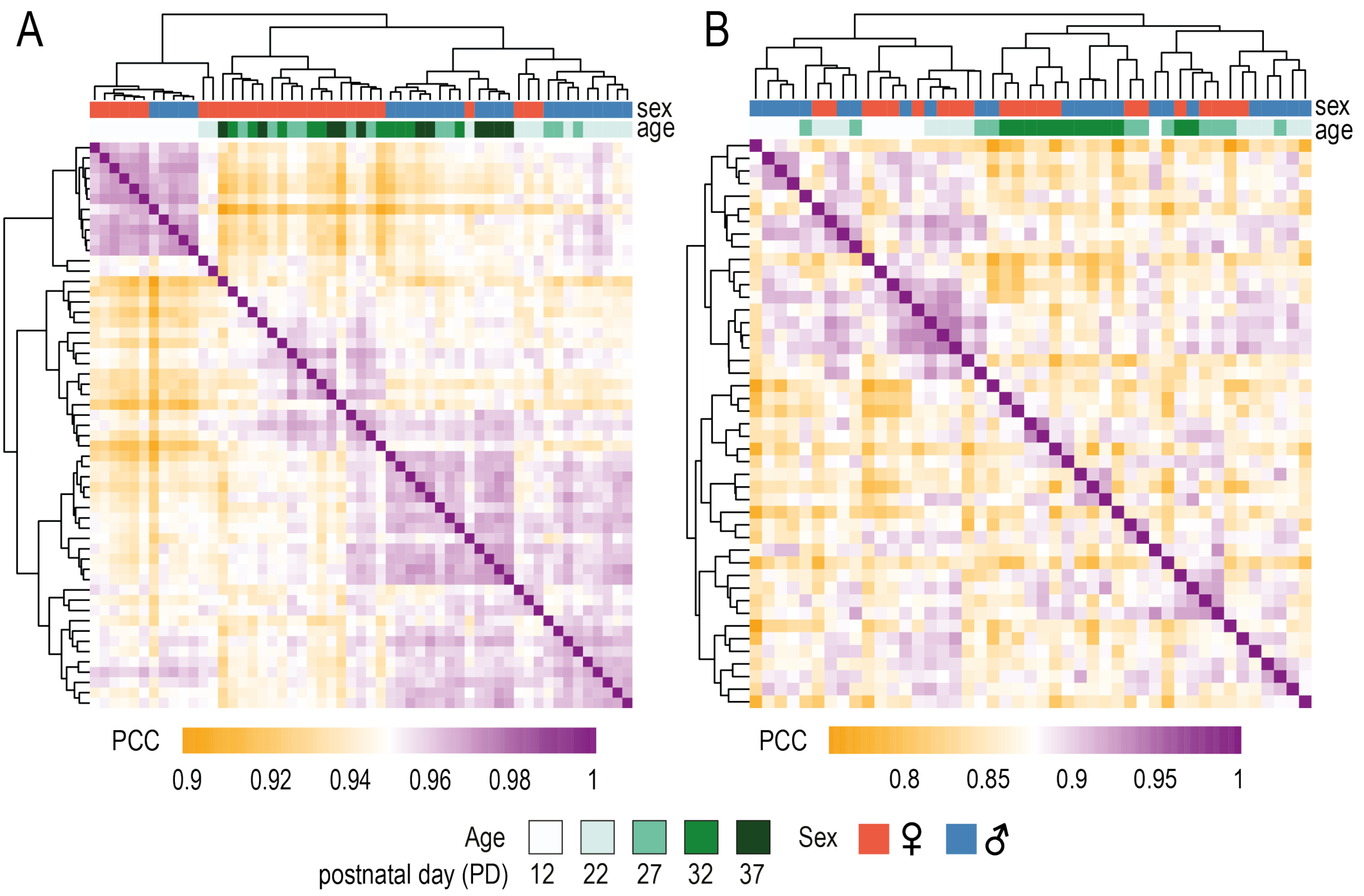


**Supplementary figure 2. Correlation heatmaps for pituitary gland samples.**
Pearson correlation between samples is calculated based on **(A)** gene expression and **(B)** miRNA expression. Hierarchical clustering is performed based on pairwise Pearson correlation coefficients (PCCs). Heatmap color intensity represents PCCs (orange: lower correlation; purple: higher correlation). Sample conditions are shown in top bars. Green shades: different ages, blue: male samples; red: female samples.

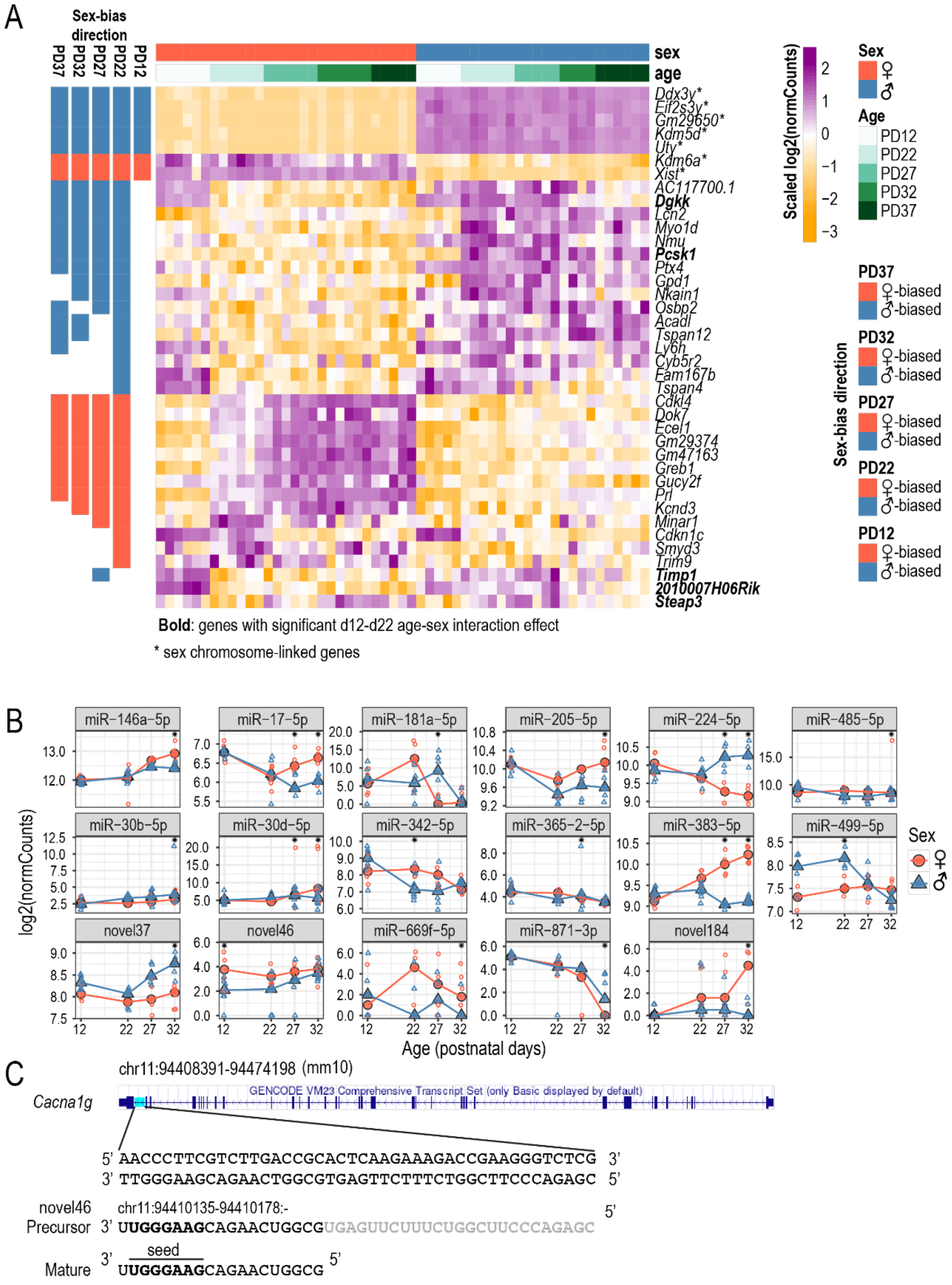
A
Sex-bias direction
PD37
PD32
PD27
PD22
PD12
sex
age
Ddx3y*
Eif2s3y*
Gm29650*
Kdm5d*
Uty*
Kdm6a*
Xist*
AC117700.1
Dgkk
Lcn2
Myo1d
Nmu
Pcsk1
Ptx4
Gpd1
Nkain1
Osbp2
Acadl
Tspan12
Ly6h
Cyb5r2
Fam167b
Tspan4
Cdkl4
Dok7
Ecel1
Gm29374
Gm47163
Greb1
Gucy2f
Prl
Kcnd3
Minar1
Cdkn1c
Smyd3
Trim9
Timp1
2010007H06Rik
Steap3
Bold: genes with significant d12-d22 age-sex interaction effect
* sex chromosome-linked genes
Scaled log2(normCounts)
Sex
♀
♂
Age
PD12
PD22
PD27
PD32
PD37
Sex-bias direction
PD37
♀-biased
♂-biased
PD32
♀-biased
♂-biased
PD27
♀-biased
♂-biased
PD22
♀-biased
♂-biased
PD12
♀-biased
♂-biased
B
miR−146a−5p
miR−17−5p
miR−181a−5p
miR−205−5p
miR−224−5p
miR−485−5p
miR−30b−5p
miR−30d−5p
miR−342−5p
miR−365−2−5p
miR−383−5p
miR−499−5p
novel37
novel46
miR−669f−5p
miR−871−3p
novel184
log2(normCounts)
Age (postnatal days)
Sex
♀
♂
C
chr11:94408391-94474198 (mm10)
Cacna1g
GENCODE VM23 Comprehensive Transcript Set (only Basic displayed by default)
5' AACCCTTCGTCTTGACCGCACTCAAGAAAGACCGAAGGGTCTCG 3'
3' TTGGGAAGCAGAACTGGCGTGAGTTCTTTCTGGCTTCCCAGAGC 5'
novel46
chr11:94410135-94410178:-
Precursor 3' UUGGGAAGCAGAACUGGCGUGAGUUCUUUCUGGCUUCCCAGAGC 5'
seed
Mature 3' UUGGGAAGCAGAACUGGCG 5'

**Supplementary figure 3. Characterization of sex-biased mRNAs and miRNAs.**

**A.** Gene expression heatmap of sex-biased genes at PD12 and PD22, and genes with significant sex-by-age effect between PD12 and PD22. Each row represents a gene and each column represents a sample. Column annotation bars indicate sample age and sex. Colours represent row-scaled log2 (normalized counts). Whether a gene is female-biased (red) or male-biased (blue) at each corresponding age is summarized by the row annotation bars on the left. Sex chromosome-linked genes are labeled with asterisks. Genes with significant sex-by-age interaction effect between PD12 and PD22 are bolded. **B.** Expression plots of all sex-biased miRNAs. Log2(normCounts) are plotted for each miRNA across ages. Large filled points represent median expression at each age and unfilled points represent each biological replicate. Red: female samples; blue: male samples. **C.** novel46 is a mirtron of *Cacna1g*. The precursor of novel46 is expressed from the last intron of *Cacna1g* (highlighted in blue). The canonical seed region (nucleotide positions 2-8) in the mature sequence of novel46 is bolded.

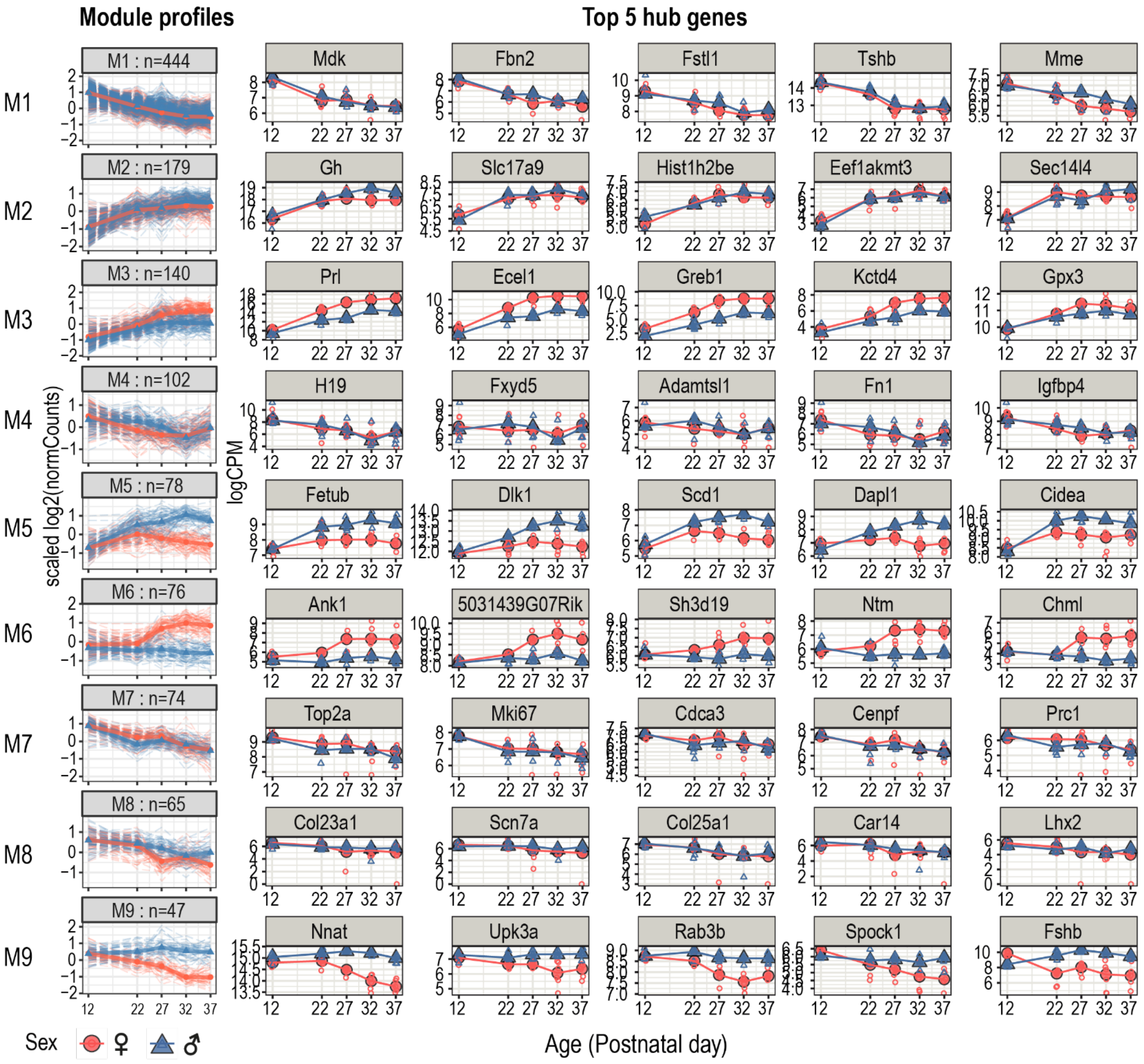


**Supplementary figure 4. Summary of gene co-expression modules.**

**Left panel.** Module expression profile (log2-transformed normalized counts (log2(normCounts)) scaled per gene across all samples) is plotted for genes within each co-expression module across profiled postnatal ages. Dotted lines: individual gene profiles, solid line: median expression profile of module genes. Number of genes within each module is labelled at the top of the plots. Red: female samples; blue: male samples. **Right panel.** Expression profile of top five hub genes for each module. log2(normCounts) is plotted across profiled postnatal ages. Large, filled points represent median expression at each age and unfilled points represent each biological replicate. Blue: male samples; red: female samples.

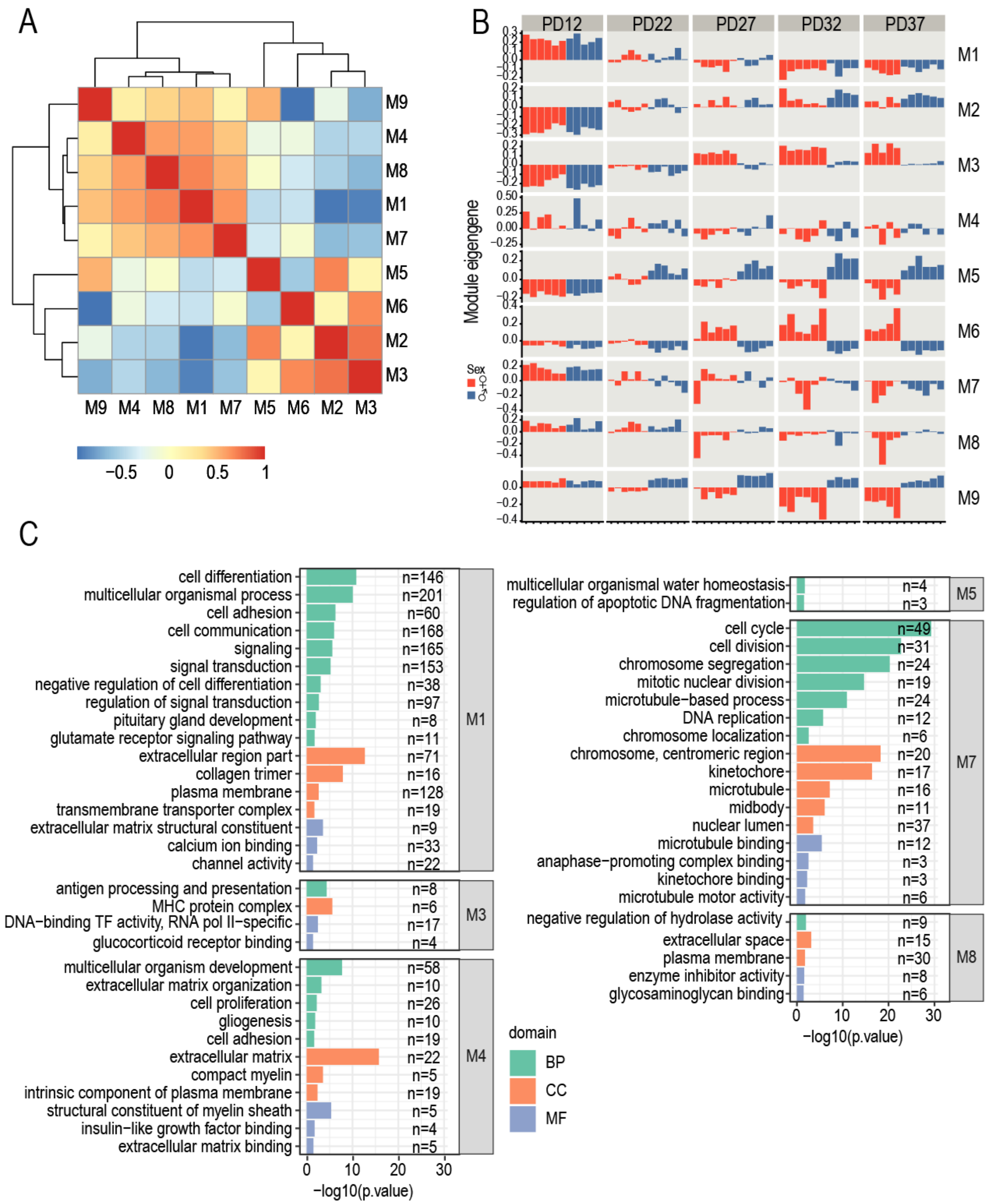


**Supplementary figure 5. Characterization of co-expression gene modules.**

**A.** Correlation heatmap based on module eigengene (first PC, obtained using "mod_summary()" function from "CEMitool"). Color scale represents pairwise Pearson correlation coefficients. **B.** Barplots showing the eigengene for each sample in each module. Samples are grouped by age and colored by sex (blue: male; red: female). **C.** Barplots showing pathway enrichment results for genes in each module. Each bar represents a pathway. Colour represents pathway categories (BP: Biological Process; CC: Cellular Component; MF: Molecular Function). Number of module genes in the pathway is labeled. X axis: -log10(*P*-value) of each pathway.

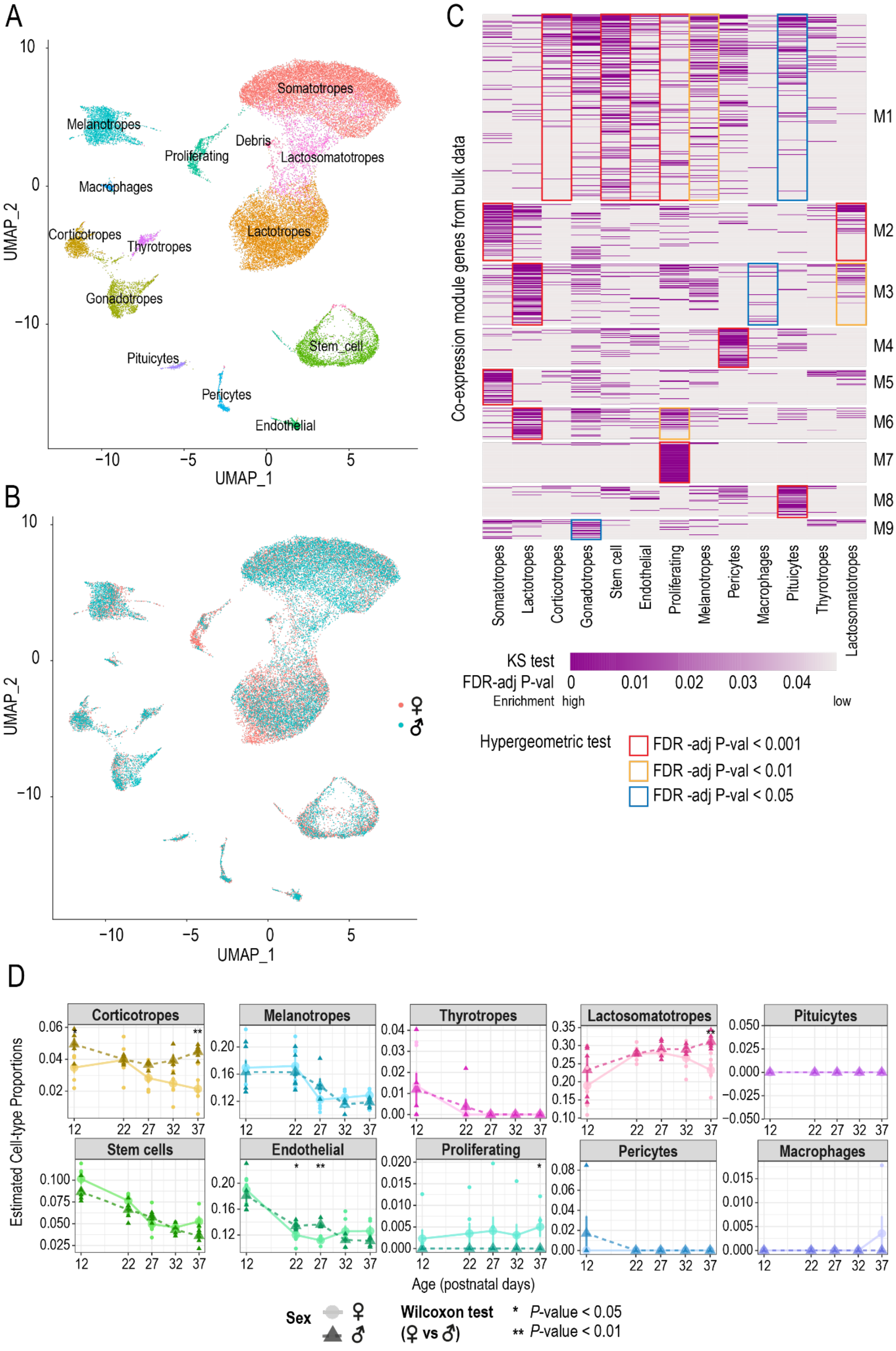

A
B
C
D
UMAP_1
UMAP_2
Somatotropes
Melanotropes
Proliferating
Debris
Lactosomatotropes
Macrophages
Corticotropes
Thyrotropes
Lactotropes
Gonadotropes
Stem_cell
Pituicytes
Pericytes
Endothelial
Co-expression module genes from bulk data
M1
M2
M3
M4
M5
M6
M7
M8
M9
KS test
FDR-adj P-val
Enrichment high
low
Hypergeometric test
FDR -adj P-val < 0.001
FDR -adj P-val < 0.01
FDR -adj P-val < 0.05
Estimated Cell-type Proportions
Corticotropes
Melanotropes
Thyrotropes
Lactosomatotropes
Pituicytes
Stem cells
Endothelial
Proliferating
Pericytes
Macrophages
Age (postnatal days)
Sex
Wilcoxon test
(♀ vs ♂)
* P-value < 0.05
** P-value < 0.01

**Supplementary figure 6. Co-expression module gene enrichment in single-nuclei RNA-seq data.**

Uniform Manifold Approximation and Projection (UMAP) dimension reduction representation of adult female and male mouse pituitary gland single-nuclei transcriptome integrated by sex from Ruf-Zamojski et al. 2021 **(A)** coloured by cell type and **(B)** coloured by sample sex**.** Samples derived from snap-frozen pituitaries were first merged between replicates for each sex (n=3/sex). Merged samples were then integrated between sexes. **C.** Enrichment heatmap of co-expression module genes within cell types. One-sided Kolmogorov-Smirnov (KS) test was performed to test for enrichment of each co-expression module gene within a given cell type compared to all other cell types. Colour gradient represents KS test FDR-adjusted $P-$value for each gene (dark purple: high enrichment; gray: low enrichment). Only genes with FDR ≤ 0.05 in at least one cell type are plotted and only FDR < 0.05 are shown. Each column represents a cell type as labelled at the bottom of the heatmap. Each row represents a co-expression module gene and the genes are grouped by the co-expression module in which the gene was identified (labelled on the right). Breaks were added in the heatmap between co-expression modules. Coloured boxes represent the level of significance based on a one-sided hypergeometric test to determine if a group of module genes with KS test FDR ≤ 0.05 was significantly enriched for a given cell type. **D.** Estimated cell-type proportions by RNA-seq deconvolution using Proportions in Admixture (WGCNA) changes across profiled ages of pituitary cell types without known sex differences in their proportions. Estimated cell-type proportions are plotted across postnatal ages along the x-axis. Large circles and triangles represent the mean cell-type proportion at each age and small circles and triangles represent each biological replicate. Lighter colour, solid line, circle points: female samples; dark colour, dotted line, triangle points: male samples. Wilcoxon test was performed to compare cell proportions between both sexes at each age (*P<0.05, **P<0.01). See **Figure 5B** for estimated proportions of cell types with known sex biases.
